# Degenerate circuits use distinct mechanisms to respond similarly to the same perturbation

**DOI:** 10.1101/2020.07.24.220442

**Authors:** D.J. Powell, E. Marder, M.P. Nusbaum

## Abstract

There is considerable flexibility embedded within neural circuits. For example, separate modulatory inputs can differently configure the same underlying circuit but these different configurations generate comparable, or degenerate, activity patterns. However, little is known about whether these mechanistically different circuits in turn exhibit degenerate responses to the same inputs. We examined this issue using the crab (*Cancer borealis*) stomatogastric nervous system, in which stimulating the modulatory projection neuron MCN1 and bath applying the neuropeptide CabPK II elicit similar gastric mill (chewing) rhythms in the stomatogastric ganglion, despite differentially configuring the same neural circuit. We showed previously that bath applying the peptide hormone CCAP or stimulating the muscle stretch-sensitive sensory neuron GPR during the MCN1-elicited gastric mill rhythm selectively prolongs the protraction or retraction phase, respectively. Here, we found that these two influences on the CabPK-rhythm elicited some unique and unexpected consequences compared to their actions on the MCN1-rhythm. For example, in contrast to its effect on the MCN1-rhythm, CCAP selectively decreased the CabPK-rhythm retraction phase and thus increased the rhythm speed, whereas the CabPK-rhythm response to stimulating GPR during the retraction phase was similar its effect on the MCN1-rhythm (i.e. prolonging retraction). Interestingly, despite the comparable GPR actions on these degenerate rhythms, the underlying synaptic mechanism was distinct. Thus, degenerate circuits do not necessarily exhibit degenerate responses to the same influence, but when they do, it can occur via different underlying mechanisms.

**Significance Statement:** Circuits generating seemingly identical behaviors are often thought to arise from identical circuit states, as that is the most parsimonious explanation. Here we take advantage of an alternate scenario wherein a well-defined circuit with known connectivity generates similar activity patterns using distinct circuit states, via known mechanisms. The same peptide hormone modulation of these distinct circuit states produced divergent activity patterns, whereas the same sensory feedback altered these circuit outputs similarly but via different synaptic pathways. The latter observation limits the insights available from comparable studies in systems lacking detailed access to the underlying circuit.

## Introduction

It is often assumed that indistinguishable neuronal activity across individuals or across time points in the same individual indicates a conserved circuit or circuit state. In many cases, however, limited access to the underlying circuit prevents confirming or disproving this assumption. Moreover, there is an alternative possibility because there are also distinct neural circuits or circuit states that produce similar output both across and within individuals (Marder and Weimann, 1992; Goldman et al., 2001; Prinz et al., 2004; Schulz et al., 2006; Saideman et al., 2007a; Beverly et al., 2011; Gutierrez et al., 2013; Rodriguez et al., 2013; Trojanowski et al., 2014; Cropper et al., 2016; Sakurai and Katz, 2017; Alonso and Marder, 2019; Wang et al., 2019; Onasch and Gjorgjieva, 2020). This latter phenomenon is termed degeneracy (Tononi et al., 1999; Edelman and Gally, 2001; Leonardo, 2005; Marder et al., 2015; Ikeda et al., 2020).

The presence of degenerate circuits raises the issue of whether such circuits respond similarly or distinctly to a consistent perturbation, such as hormonal modulation or a sensory input. The fact that degenerate circuits are distinct suggests that each such circuit could respond differently to the same perturbation, as shown recently by Wang et al. (2019). We are addressing this issue in the stomatogastric ganglion (STG) of the crab *Cancer borealis*, which contains the well-characterized gastric mill (chewing) and pyloric (pumping and filtering of chewed food) circuits (Marder and Bucher, 2007; Daur et al., 2016; Nusbaum et al., 2017).

There are at least several different versions of the gastric mill rhythm in *C. borealis* (Blitz et al., 1999; Beenhakker and Nusbaum, 2004; Blitz et al., 2004; Blitz et al., 2008; White and Nusbaum, 2011; Blitz et al., 2019). Additionally, there is gastric mill circuit degeneracy, as the gastric mill rhythms driven by stimulating modulatory commissural neuron 1 (MCN1) and bath-applying CabPK (*C. borealis* pyrokinin) peptide are comparable, despite being generated by distinct cellular and synaptic mechanisms (Coleman and Nusbaum, 1994; Saideman et al., 2007a; Stein et al., 2007; Rodriguez et al., 2013).

To investigate whether the MCN1- and CabPK-driven gastric mill rhythms respond degenerately to identical extrinsic inputs, we chose two inputs common to all nervous systems, hormonal modulation and sensory feedback. Previous work established that the peptide hormone CCAP (crustacean cardioactive peptide) slows the MCN1-gastric mill rhythm by selectively prolonging protraction (Kirby and Nusbaum, 2007; DeLong et al., 2009b), while stimulating the muscle stretch-sensitive sensory Gastropyloric Receptor neuron (GPR) slows the MCN1-rhythm by selectively prolonging retraction (Beenhakker et al., 2005; DeLong et al., 2009a). Here, we studied the CabPK-gastric mill rhythm response to CCAP and GPR and, to ensure continuity, in the same experiments again assessed the MCN1-gastric mill rhythm response to both perturbations. Our results demonstrate that similar neural output generated by distinct circuit states can in turn produce either divergent or convergent output in response to a shared input. Additionally, such convergent output does not necessarily result from the presence of a conserved synaptic mechanism between the different circuit states.

## Materials and Methods

### Animals

Male Jonah crabs (*C. borealis*) were purchased from Commercial Lobster (Boston, MA) and maintained in filtered, circulating artificial sea water (10-12° C) on a 12hr/12hr, light/dark cycle, typically 1 week prior to dissection. Crabs were anesthetized for 30 min by packing in ice prior to dissection. Dissections were performed as previously described (Gutierrez and Grashow, 2009). Each dissected STNS was pinned down in a Petri dish with a silicone elastomer-lined (Sylgard 184: Dow Corning) bottom.

### Peptides & Solutions

Dissected nervous systems were maintained in *C. borealis* saline (440 mM NaCl, 11 mM KCl, 26 mM MgCl_2_, 13 mM CaCl_2_, 11 mM Trizma base, 5 mM maleic acid, pH 7.45 at 21° C) throughout the dissection and experiment. Saline or peptide solution was continuously superfused (10 ml/min) at 10-12°C across the preparation for the duration of each experiment. Saline temperature was maintained within this narrow range because these animals are poikilotherms and so neuronal activity is affected by temperature (Tang et al., 2010; Stadele et al., 2015; Powell et al., 2020).

Peptide solutions were superfused across the STNS at 10-12° C, either individually or in combination (i.e. CabPK + CCAP). *Cancer borealis* pyrokinin peptide II (CabPK II: SGGFAFSPRL-NH_2_ (Saideman et al., 2007b); GenScript) was dissolved into DDI-H_2_O to form a stock solution (10^−3^ M). This stock solution was then diluted into *C. borealis* saline to the concentration used for application (10^−6^ M). Crustacean cardioactive peptide (CCAP: PFCNAFTGC-NH_2_; Bachem) was similarly dissolved into DDI-H_2_O to create a stock solution (10^−3^ M), which was further diluted in *C. borealis* saline (10^−6^ M) for all experiments. For experiments where these two peptides were co-applied, the peptides were diluted into the same solution. To avoid the possibility that there were sequence-dependent effects of this experimental manipulation, we counterbalanced the sequence of peptide applications in different preparations.

For experiments in which the Lateral Gastric (LG) neuron was isolated from the other STG neurons, the STNS was superfused with *C. borealis* saline containing picrotoxin (PTX; 10^−5^ M; Sigma), which in this system blocks all ionotropic inhibitory glutamatergic receptors, which mediate most inhibitory synapses in the STG (Bidaut, 1980), and all STG synapses onto LG.

### Electrophysiology

In all experiments, the CoGs were removed from the isolated STNS by bisecting the superior (*sons*)- and inferior oesophageal nerves (*ions*), to eliminate CoG projection neuron activity from influencing the STG circuits and prevent feedback from STG and GPR neurons to these projection neurons (**Fig. 1A**). The CoG projection neuron MCN1 was selectively stimulated via extracellular *ion* stimulation (Bartos and Nusbaum, 1997). STNS neuron action potentials were recorded extracellularly from isolated sections of nerves using pairs of stainless steel pin electrodes (reference and recording) and amplified via model 3500 AC amplifiers (A-M Systems). To this end, sections of nerve were isolated from the saline bath either by 1) building a Vaseline barrier around a section of each nerve, or 2) sealing both electrode pins to a length of nerve with Vaseline such that the only conduction between the electrode tips was through the nerve. For most STG peripheral nerves (*lgn, dgn, mvn, lvn, pdn*), recordings were obtained using barriers, while the sealed approach was used for the *ions* and *gpns*, to enable lower voltage stimulation of MCN1 or GPR, respectively (**Fig. 1A**). Recorded neurons were identified based on nerve recordings and activity patterns (Weimann et al., 1991).

**Figure 1.**
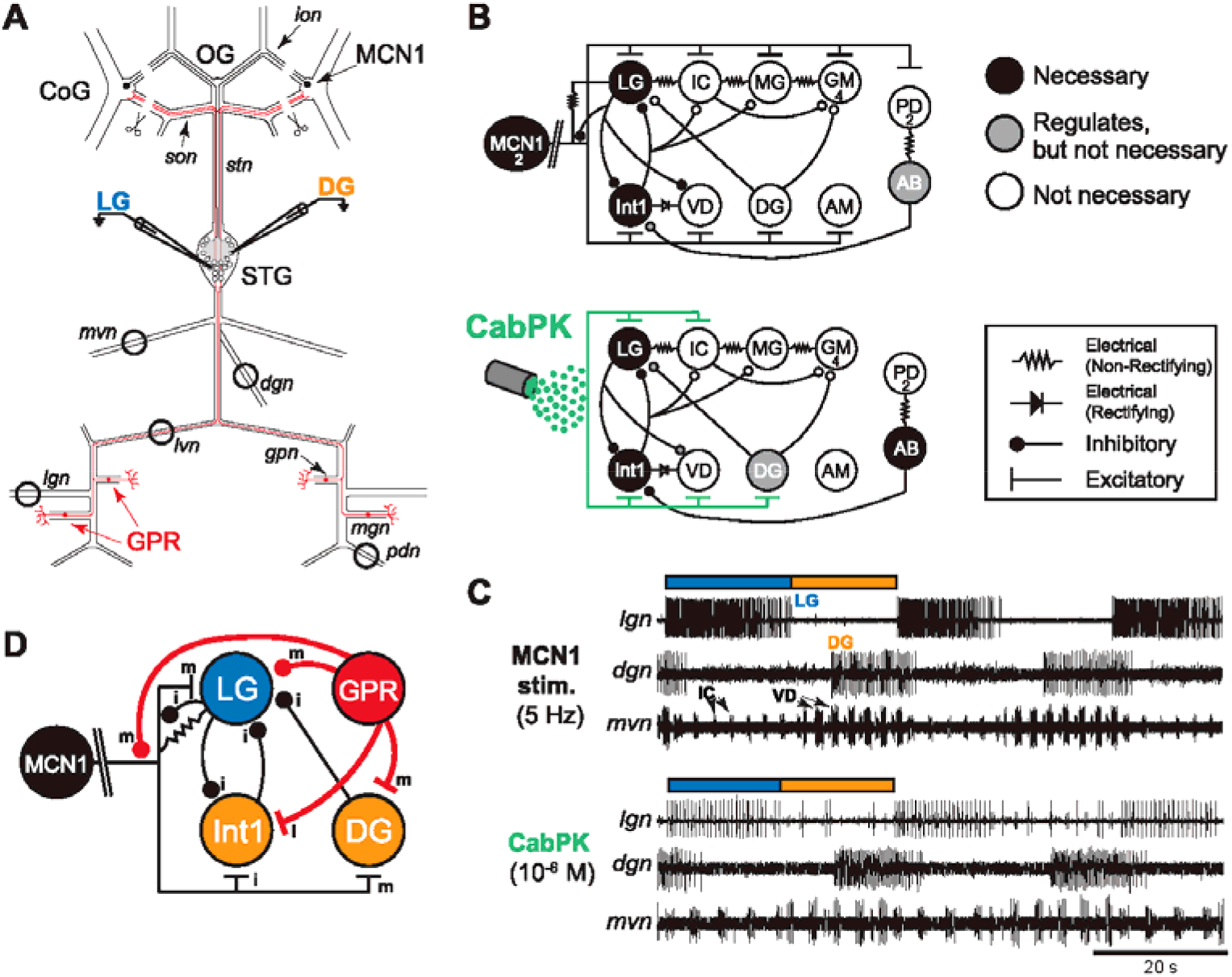
Distinct circuit states can generate similar gastric mill rhythms. ***A***, Schematic of the *C. borealis* STNS. Black circles surrounding nerves indicate placement of Vaseline barriers for extracellular nerve recordings. Scissor symbols indicate where *ions* and *sons* were bisected for these experiments. Microelectrodes represent intracellular recordings of the gastric mill neurons LG and DG. Red lines represent the axon projections of GPR. Abbreviations: Ganglia-CoG, commissural ganglion, OG; oesophageal ganglion; STG, stomatogastric ganglion. Neurons: DG, dorsal gastric; GPR, gastropyloric receptor; LG, lateral gastric; MCN1, modulatory commissural neuron 1. Nerves: *dgn*, dorsal gastric nerve; *gpn*, gastropyloric nerve; *lgn*, lateral gastric nerve; *lvn*, lateral ventricular nerve; *ion*, inferior oesophageal nerve; *mgn*, medial gastric nerve; *mvn*, medial ventricular nerve; *pdn*, pyloric dilator nerve; *son*, superior oesophageal nerve; *stn*, stomatogastric nerve. ***B***, Circuit diagram schematics for the MCN1- (top) and CabPK- (bottom) gastric mill rhythms. Top row, gastric mill protractor neurons; bottom row, gastric mill retractor neurons; AB/PD, pyloric pacemaker neurons. Color of each neuron indicates its role in rhythm generation, as indicated in the legend to the right of the circuit schematics. The double backslash lines across the MCN1 axon represent additional distance between the MCN1 soma in the CoG and its axon terminals in the STG. All indicated synapses occur in the STG. The numbers inside some circles indicate the copy number for neurons present as more than one copy per STNS. Note that applied CabPK does not synapse onto neurons but does excite all indicated neuron targets. Modified from: Saideman et al. (2007a). ***C***, Recordings of the MCN1- (top) and CabPK-(bottom) elicited gastric mill rhythms. Blue and orange bars indicate protraction and retraction durations, respectively. STG neurons/labels are colored based on the phase in which they are active. Both gastric mill rhythms were recorded in the same preparation. ***D***, Simplified circuit schematic of the GPR synapses onto gastric mill rhythm generator neurons LG, Int1, and MCN1, plus the retractor neuron DG. Neuron colors are consistent with (***C***). MCN1 and GPR are colored black and red, respectively, in all figures. Note that the LG synapse onto MCN1 occurs within the STG. Abbreviations: i, ionotropic; m, metabotropic.

The STG was desheathed dorsally and neuron somata were impaled using glass microelectrodes (ID: 0.86 mm; OD: 1.5 mm; 5-20 MΩ; Sutter Instrument). Microelectrodes were backfilled with a solution containing 10 mM MgCl_2_, 400 mM potassium gluconate, 10 mM HEPES buffer, 15 mM Na_2_SO_4_, and 20 mM NaCl (Hooper et al., 2015). Electrodes were pulled on a P-97 micropipette puller (Sutter Instrument Co.). To visualize neurons for impalement, white light was directed through a dark-field condenser (Nikon) to the STG from below the Petri dish. Intracellular membrane voltage was recorded using a model 900A amplifier (Axon Instruments/Molecular Devices). In some cases, intracellular and extracellular recordings were further amplified and filtered using model 410 amplifiers (Brownlee Precision). Analog signals were digitized (10-20 kHz) using either a Digidata 1440 (Axon Instruments/Molecular Devices) or a Micro 1401 A/D board (Cambridge Electronic Design). Data were recorded using both pClamp data acquisition software (Axon Instruments/Molecular Devices, version 10.5) and Spike2 (version 6; Cambridge Electronic Design) respective to the A-D converter used. For a handful of experiments, data were simultaneously recorded on an Everest model chart recorder (AstroNova) and computer.

MCN1 stimulation (3-6 Hz) to drive the gastric mill rhythm was performed by simultaneous stimulation of both *ions* using the same stimulation frequency. MCN1 stimulation was continuous, using constant inter-stimulus intervals, with manual initiation and cessation using a model 3800 stimulator (A-M Systems) via model 3820 stimulus isolation units (A-M Systems). Each stimulus duration was 0.3 ms at just supra-threshold voltage. Stimulation of the *ions* at this relatively low frequency/voltage range ensures that MCN1 is selectively stimulated (Coleman et al., 1995; Bartos and Nusbaum, 1997). Threshold for MCN1 stimulation was determined by monitoring the electrical EPSPs (eEPSPs) produced by MCN1 in intracellular recordings of the LG neuron (Coleman et al., 1995).

GPR stimulation (*gpn:* stim. freq., 6-8 Hz; individual stim. durations, 0.3 ms) was delivered at just supra-threshold voltage using a model 3800 stimulator (A-M Systems) via model 3820 stimulus isolation units (A-M Systems). GPR activation of Dorsal Gastric (DG; *dgn*) neuron bursting was used to determine GPR stimulation threshold (Katz and Harris-Warrick, 1989; Kiehn and Harris-Warrick, 1992). The *dgn* contains the axons of the DG, AGR (muscle sensory neuron) and Gastric Mill (GM) neurons. In *dgn* recordings, DG activity is readily distinguished from AGR and GM activity based on activity pattern and action potential amplitude (Smarandache and Stein, 2007). Additionally, neither GM nor AGR is activated by GPR (Blitz et al., 2004). Each gastric mill rhythm-timed GPR stimulation was performed manually during either the protraction or retraction phase. These stimulation durations were time-matched to the control protraction or retraction phase duration.

In experiments in which DG was hyperpolarized, a single bridge balanced electrode was used to inject hyperpolarizing current. For all experiments where LG was isolated using PTX (10^−5^ M) saline, we used two-electrode current clamp (TECC) to manipulate the LG membrane potential. To depolarize LG sufficiently in these experiments to generate a modest firing rate, constant amplitude and duration depolarizing steps (1-4 nA) were delivered to LG through an intracellular electrode either manually or using software protocols available in pClamp. The magnitude of the current step depended on the LG input resistance (10-20 MΩ). Neurons with an input resistance below 10 MΩ were not used.

### Dynamic clamp

We used the dynamic clamp to inject an artificial version of the modulator activated, voltage-sensitive inward current (I_MI_) into the LG neuron (Sharp et al., 1993a, b; Bartos et al., 1999; Swensen and Marder, 2001; Goaillard and Marder, 2006; DeLong et al., 2009b; Rodriguez et al., 2013). In particular, we used the dynamic clamp to inject an artificial CCAP-activated I_MI_ (I_MI-CCAP_) (DeLong et al., 2009b). In these experiments, the LG neuron membrane voltage (V_m_) was recorded in TECC or in discontinuous current clamp (DCC; sampling rate 3 kHz). We developed a custom module for dynamic clamp software (RTXI, www.rtxi.org). This module continuously monitors V_m_ and calculates a current based on equations describing I_MI_ (I_dyn_) to inject into LG. The magnitude of I_MI_ is determined by a variety of free parameters, including the: I_MI_ reversal potential (E_rev_), I_MI-CCAP_ conductance (g_dyn_), half-activation voltage (V_1/2_) of I_MI_, and slope of the half-activation curve (*k*_X_). *X*represents either the activation (*m*) or inactivation (*h*) gating functions. The membrane time constants above and below V_1/2_ (τ_Hi_ and τ_Lo_, respectively) also shape activation of the injected current. Because V_1/2_ can be different for the *m*_∞_ and *h_∞_* functions, it is generalized as V_X_. The injected current is based on real-time calculations and updated each time step (0.04 ms) to match new values of V_m_ and was computed using the following equations:

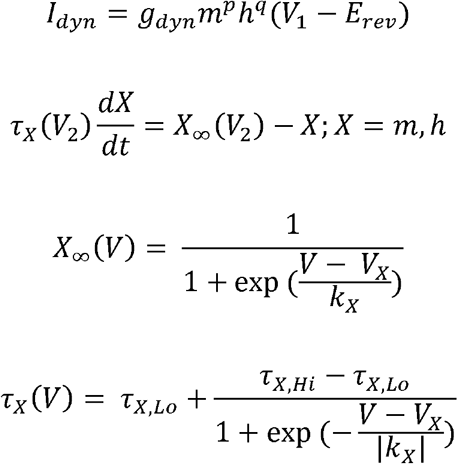

The values for free parameters used here were based on voltage clamp data (Golowasch and Marder, 1992; Swensen and Marder, 2000, 2001; DeLong et al., 2009b; Rodriguez et al., 2013). The free parameters were set to the following values: E_rev_ = +10 mV, V_1/2_ = −45 mV, k_m_ = −5 mV, τ_m,Hi_ = 100 ms, τ_m,Lo_ = 50 ms. I_MI_ has a voltage dependent activation (*m*), and the integer power of activation (*p*) was set to 1 (Golowasch and Marder, 1992). I_MI_ does not inactivate, so the inactivation variable *h* and associated exponent *q* were set to 0. Thus, parameters for k_h_, τ_h,Hi_, and τ_h,Lo_ were not set or used for dynamic clamp calculations. For these experiments we used a range of g_MI_ values (5-120 nS) to recapitulate the effect of CCAP bath application during the CabPK rhythm, as I_MI-CCAP_ was not measured in each LG neuron. As each LG neuron will have unique intrinsic properties (Schulz et al., 2006; Schulz et al., 2007) and leak current from impalement (R_in_ = 10-20 MΩ), a range of values was needed. In all dynamic clamp experiments, the maximum positive current injected never exceeded 3 nA and on average was 1-2 nA. This range of maximal current is consistent with voltage clamp data of I_MI-CCAP_ (DeLong et al., 2009b; Rodriguez et al., 2013). Using this range of values, we selected values of g_MI_ that mimicked the effect of CCAP on the LG neuron (see **Results**). I_MI-CCAP_ dynamic clamp module is available at https://github.com/dpo3205/Powell-Marder-Nusbaum-2020.

### Data analysis

Data analysis was performed using both Spike2 (version 7.01; Cambridge Electronic Design) and MATLAB (MathWorks) using custom scripts that are available at: https://github.com/dpo3205/Powell-Marder-Nusbaum-2020. For all gastric mill rhythm experiments, 2-4 min of continuous recordings were used to generate means and standard deviations (mean ± SD) for each experiment. For each condition, individual extracellularly recorded bursts from each neuron were analyzed using customized Spike2 scripts to determine burst duration, burst frequency, the start and stop times for each burst, within burst spike frequency, and spike count per burst.

Burst onset and offset were defined by the first and last spike in a burst. Bursts were defined by an interspike interval (ISI) threshold (~0.8 s) where spikes with a longer ISI were excluded. Spikes that are close to but not part of an LG burst typically are pyloric rhythm-timed and occur one or more pyloric cycles adjacent to the LG burst. Therefore, we used an ISI threshold that was less than a single pyloric cycle in a given preparation. With intracellular LG recordings, bursts were further defined to include all action potentials occurring during an LG burst plateau. Most experiments included an intracellular LG recording (*N* = 31/45). Protraction duration was defined as the LG burst duration, while retraction duration was the LG interburst interval. Cycle period was calculated as the sum of the protraction and retraction durations. Within burst spike frequency was calculated by dividing the number of spikes in a burst minus one, by the burst duration.

The pyloric cycle frequency was determined as one divided by the pyloric cycle period (i.e. duration from the onset of two consecutive PD neuron (*pdn*) bursts). Mean pyloric cycle frequency was calculated from the instantaneous PD cycle frequency during an ongoing gastric mill rhythm. To investigate if the CCAP-selective decrease in CabPK retraction duration was due to a CCAP driven change in pyloric frequency, we calculated the ratio of pyloric cycles to gastric mill cycles, and the pyloric cycle frequency as a function of neuromodulatory drive (CabPK and CCAP). For this analysis, we excluded experiments where CCAP did not affect the CabPK-gastric mill rhythm [*N* = 6/45 (all experiments) and *N* = 1/9 (pyloric frequency analysis)].

For all descriptive statistics, we report mean ± SD. All statistics can be found in either the Figure Legends or Tables. Metadata for all experiments are in an Excel file on the same GitHub link as the analysis scripts. Statistical analyses were carried out using both MATLAB (MathWorks) and SPSS (IBM). Figures were made using Clampfit 10.5 (Axon Instruments/Molecular Devices), MATLAB (MathWorks), and Adobe Illustrator (CC 2018; Adobe).

## Results

### The *C. borealis* gastric mill rhythms

The dorsally-located stomatogastric nervous system (STNS), which wraps around the decapod crustacean foregut, is composed of four ganglia that regulate, generate, and coordinate rhythmic motor patterns which are named for the stomach compartment whose movement they control (Marder and Bucher, 2007). These ganglia include the STG, oesophageal ganglion (OG) and the paired commissural ganglia (CoGs) (**Fig. 1A**). The foregut is composed of the oesophagus plus a three-compartment stomach. These compartments include, from anterior to posterior, the cardiac sac (food storage), gastric mill (chewing) and pylorus (pumping/filtering of chewed food). In particular, rhythmic contractions of gastric mill muscles produce the relatively slow (~10-20 sec/cycle) gastric mill chewing patterns, while faster (~1 sec/cycle) rhythmic contractions of pyloric muscles drive the pyloric filtering patterns. These two patterns are generated by overlapping neural circuits in the STG (Marder and Bucher, 2007; Nusbaum et al., 2017). Each phase (protraction, retraction) of the gastric mill rhythm corresponds to the position of the three gastric mill ossicles or ‘teeth’, each of which is tethered at one end to the internal stomach wall in the gastric mill compartment (Heinzel, 1988; Heinzel et al., 1993; Diehl et al., 2013). During protraction, muscle contractions move the teeth together to generate a chew, while the teeth are moved away from one another by a distinct set of muscle contractions during retraction.

In *C. borealis*, 22/26 STG neurons (20 motor neurons and 2 interneurons) contribute to the gastric mill and/or pyloric circuit. One interneuron (AB) is a rhythmically bursting oscillator that drives the pyloric rhythm and either regulates or is a necessary component of the gastric mill rhythm generator (**Fig. 1B**). The core of gastric mill rhythm generator includes a half-center oscillator (i.e. reciprocally inhibitory pair) composed of the retraction phase neuron Interneuron 1 (Int1) and the LG protractor motor neuron. This half-center is pivotal to all characterized gastric mill rhythms (Coleman et al., 1995; Nadim et al., 1998; Saideman et al., 2007a; White and Nusbaum, 2011).

The bi-phasic gastric mill rhythm includes bursting activity from four types of protractor motor neurons (LG, MG, GM, IC) and four retractor phase neurons (Int1; motor neurons: DG, AM, VD) (**Fig. 1B**). All these neurons occur as single copies, except for GM which is present as four apparently equivalent copies. Each phase of the gastric mill rhythm is driven primarily by the synaptic actions of the half-center neurons LG and Int1 and those of the activating system (e.g. MCN1, CabPK) (**Fig. 1B**) (Saideman et al., 2007a,b; Rodriguez et al., 2013; Nusbaum et al., 2017). Therefore, we used LG burst (protraction) and interburst interval (retraction) durations to draw comparisons between the CabPK- and MCN1(3-6 Hz)-gastric mill rhythms (**Fig. 1C**).

The gastric mill rhythm is an episodic motor pattern *in vivo* and in the isolated STNS, because the projection neurons that drive this rhythm require extrinsic drive from sensory neurons and inputs from other regions of the CNS (Beenhakker et al., 2004; Blitz et al., 2004; Christie et al., 2004; Blitz et al., 2008; Diehl et al., 2013). Most of these projection neurons are located in the CoGs (Coleman et al., 1992; Coleman et al., 1994). Several of the distinct versions of the gastric mill rhythm include the activity of MCN1, which is present as a single copy in each CoG (Coleman and Nusbaum, 1994). Selective, tonic MCN1 stimulation is sufficient to drive the gastric mill rhythm in the isolated STNS, even with the CoGs removed (Bartos and Nusbaum, 1997).

### The degenerate MCN1- and CabPK-elicited gastric mill rhythms

MCN1, which co-localizes GABA, proctolin, and CabTRP Ia, drives the gastric mill rhythm generator via a GABA-mediated fast excitation of Int1 and a slow, CabTRP Ia-mediated excitation of LG, as well as driving LG via an electrical synapse (**Fig. 1B,D**) (Coleman et al., 1995; Blitz et al., 1999; Stein et al., 2007). MCN1 excites the remaining gastric mill neurons, as well as AB, by both peptide cotransmitters (Stein et al., 2007). AB activity regulates, but is not necessary for MCN1-gastric mill rhythm generation (Bartos et al., 1999) (**Fig. 1B**). The metabotropic MCN1 action on LG results from CabTRP Ia activation of I_MI_, a modulator-activated, voltage-sensitive inward current (Swensen and Marder, 2000; DeLong et al., 2009b). In turn, LG regulates MCN1 transmitter release via an ionotropic inhibitory synapse onto the STG terminals of MCN1 (MCN1_STG_) (**Fig. 1B,D**) (Coleman and Nusbaum, 1994). This inhibition from LG limits MCN1_STG_ cotransmitter release to the gastric mill retraction phase and is pivotal to gastric mill rhythm generation, thus defining MCN1 as both a driver and rhythm generator component of this gastric mill rhythm (Coleman et al., 1995; DeLong et al., 2009b).

In brief, during the MCN1-gastric mill retraction phase, MCN1 slow excitation of LG enables LG to gradually escape from Int1 inhibition and become active, inhibiting both Int1 and MCN1 STG, thereby terminating retraction and comimencing protraction (Coleman and Nusbaum, 1994; DeLong et al., 2009b). Despite inhibiting MCN1 transmitter release, LG activity persists for several seconds due to the slow decay of I_MI_ and the continuing presence of electrical EPSPs (eEPSPs) from MCN1 (Coleman et al., 1995). When I_MI_ decays sufficiently, LG activity terminates, enabling both MCN1 transmitter release and Int1 activity (retraction phase) to resume (Coleman et al., 1995; DeLong et al., 2009b).

CabPK application drives a gastric mill rhythm similar to the MCN1-rhythm by also exciting LG and Int1 (**Fig. 1B,C**) (Saideman et al., 2007a,b; Rodriguez et al., 2013). However, two CabPK-activated ionic currents (I_MI_, I_Trans-LTS_) are necessary for LG neuron burst generation during this rhythm (Rodriguez et al., 2013), in contrast to the single current (I_MI_) activated in LG during the MCN1-rhythm. The way in which these two CabPK-activated currents operate during the CabPK-rhythm results in AB neuron activity being necessary for rhythm generation (**Fig. 1B**). MCN1 does not contain CabPK, nor is it activated during the CabPK-gastric mill rhythm (Saideman et al., 2007b).

The gastric mill rhythms driven by dual MCN1 stimulation (3-6 Hz ea.) and CabPK (10^−6^ M) application are comparable (**Fig. 1C**), despite these inputs configuring different gastric mill circuit states that enable rhythm generation via different mechanisms (**Fig. 1B**) (Saideman et al., 2007a; Rodriguez et al., 2013). Their similarity is evident by comparing several gastric mill rhythm parameters, as well as metrics of LG neuron excitability and duty cycle within the same preparations (**Table 1**). These degenerate gastric mill circuits provided the opportunity to determine if their distinct underlying circuit mechanisms enabled them to respond differently to the same extrinsic input. To this end, we compared their responses, in the same preparations, to either the peptide hormone CCAP or the sensory feedback neuron GPR (**Fig. 1D**).

**Table 1.** Comparison of CabPK- and MCN1(3-6 Hz)-gastric mill rhythms. All values are mean ± S.D. Prot. = protraction; Ret. = retraction; Cycle Period = gastric mill rhythm cycle period; LG spike freq. = within burst spike frequency; LG Duty Cycle = fraction of the gastric mill rhythm cycle period occupied by the LG burst. All comparisons performed using paired *t*-tests with α = 0.05, *N* = 6.

|  | Prot.<br>(s) | Ret.<br>(s) | Cycle Period<br>(s) | LG<br>#Spikes/Burst | LG Spike<br>Freq. (Hz) | LG Duty<br>Cycle |
| --- | --- | --- | --- | --- | --- | --- |
| CabPK | $3.86 \pm 1.8$ | $9.21 \pm 1.5$ | $13.07 \pm 2.2$ | $17.2 \pm 6.4$ | $4.68 \pm 0.9$ | $0.30 \pm 0.09$ |
| MCN1 | $4.70 \pm 2.4$ | $11.48 \pm 3.4$ | $16.20 \pm 4.3$ | $20.4 \pm 8.0$ | $4.52 \pm 0.8$ | $0.29 \pm 0.09$ |
| *p*-values | 0.093 | 0.153 | 0.109 | 0.253 | 0.666 | 0.789 |

### CCAP selectively decreases the CabPK-gastric mill rhythm retraction phase

CCAP, which reaches the STG only as a circulating hormone, excites several gastric mill and pyloric circuit neurons, including LG, Int1, and AB, when applied to the isolated STG (Weimann et al., 1997; Kirby and Nusbaum, 2007; Garcia et al., 2015). Like CabPK, CCAP activates I_MI_ in at least several of its STG targets, including LG (Swensen and Marder, 2000, 2001; DeLong et al., 2009b). CCAP-activated I_MI_ in LG slows the MCN1-gastric mill rhythm by selectively prolonging protraction (Kirby and Nusbaum, 2007; DeLong et al., 2009b). This earlier work examined the CCAP influence on the gastric mill rhythm driven by higher frequency MCN1 stimulation (10-15 Hz) than that used in the present study (3-6 Hz). Thus, to ensure that CCAP influenced the MCN1-gastric mill rhythm driven by this lower MCN1 firing frequency, and to compare directly the CCAP actions on the MCN1(3-6 Hz)- and CabPK (10^−6^ M)-elicited gastric mill rhythms, we studied the influence of CCAP (10^−6^ M) on these latter two rhythms in the same preparations.

Co-application of CCAP and CabPK decreased the CabPK-gastric mill rhythm cycle period (~11%; *p* = 0.015) by selectively decreasing the retraction phase duration (~15%; *p* = 0.018) (**Table 2**), compared to the gastric mill rhythm produced by CabPK alone. It did not alter the number of LG spikes per burst (*p* = 0.367), but it did increase the LG intraburst spike frequency (~12%; *p* = 0.0006) (**Table 2**). Although CCAP selectively decreased the retraction duration of the CabPK-gastric mill rhythm, it did not alter the fraction of the cycle (i.e. duty cycle) occupied by the LG burst (*p* = 0.39) (**Table 2**). In 6 of 45 preparations (13%), CCAP did not decrease either retraction duration or cycle period.

**Table 2.** Comparison of CabPK-elicited gastric mill rhythm parameters in the absence and presence of CCAP (10^−6^ M). All values are mean ± SD. All column definitions same as Table 1. All comparisons performed using paired *t*-tests with α = 0.05, *N* = 18. Each bolded p-value indicates a significant difference between the two compared parameters.

|  | Prot. (s) | Ret. (s) | Cycle<br>Period (s) | LG<br>#Spikes/Burst | LG Spike<br>Freq. (Hz) | LG Duty<br>Cycle |
| --- | --- | --- | --- | --- | --- | --- |
| CabPK | 5.31 ± 2.1 | 8.54 ± 2.4 | 13.84 ± 3.5 | 25.78 ± 12.7 | 4.90 ± 1.2 | 0.38 ± 0.1 |
| CabPK+CCAP | 5.04 ± 2.5 | 7.23 ± 2.5 | 12.27 ± 4.3 | 27.41 ± 14.5 | 5.49 ± 1.2 | 0.40 ± 0.1 |
| *p*-values | 0.46 | **0.018** | **0.015** | 0.3668 | **0.0006** | 0.39 |

### CabPK- and MCN1-gastric mill rhythms diverge in the presence of CCAP

**Figure 2** shows that, in the same preparation, CCAP application selectively decreased the CabPK-rhythm retraction duration (**Fig. 2A**) resulting in a faster CabPK-rhythm, while it selectively prolonged the MCN1(3-6 Hz)-rhythm protraction phase (**Fig. 2B**). These effects were consistent across experiments (**Fig. 2C,D)**. The CCAP influence on the MCN1(3-6 Hz)-gastric mill rhythm phase durations was consistent with that reported previously during the MCN1(10-15 Hz)-rhythm (Kirby and Nusbaum, 2007; DeLong et al., 2009b). However, CCAP did not increase the MCN1(3-6 Hz)-gastric mill rhythm cycle period (*p* = 0.37; repeated paired *t*-test with a Bonferroni correction α = 0.0167, *N* = 6), which is increased by CCAP during the MCN1(10-15 Hz)-gastric mill rhythm (Kirby and Nusbaum, 2007). Consistent with the larger data set reported in **Table 2** (*N* = 18), in experiments where CCAP was applied during both the MCN1(3-6 Hz)- and CabPK-gastric mill rhythms it decreased the CabPK-rhythm cycle period (*p* = 0.014; repeated paired *t*-test with a Bonferroni correction α = 0.0167, *N* = 6). There was no difference between the cycle period of the MCN1- and CabPK-rhythms in normal saline (*p* = 0.11; repeated paired *t*-test with a Bonferroni correction α = 0.0167, *N* = 6).

**Figure 2.**
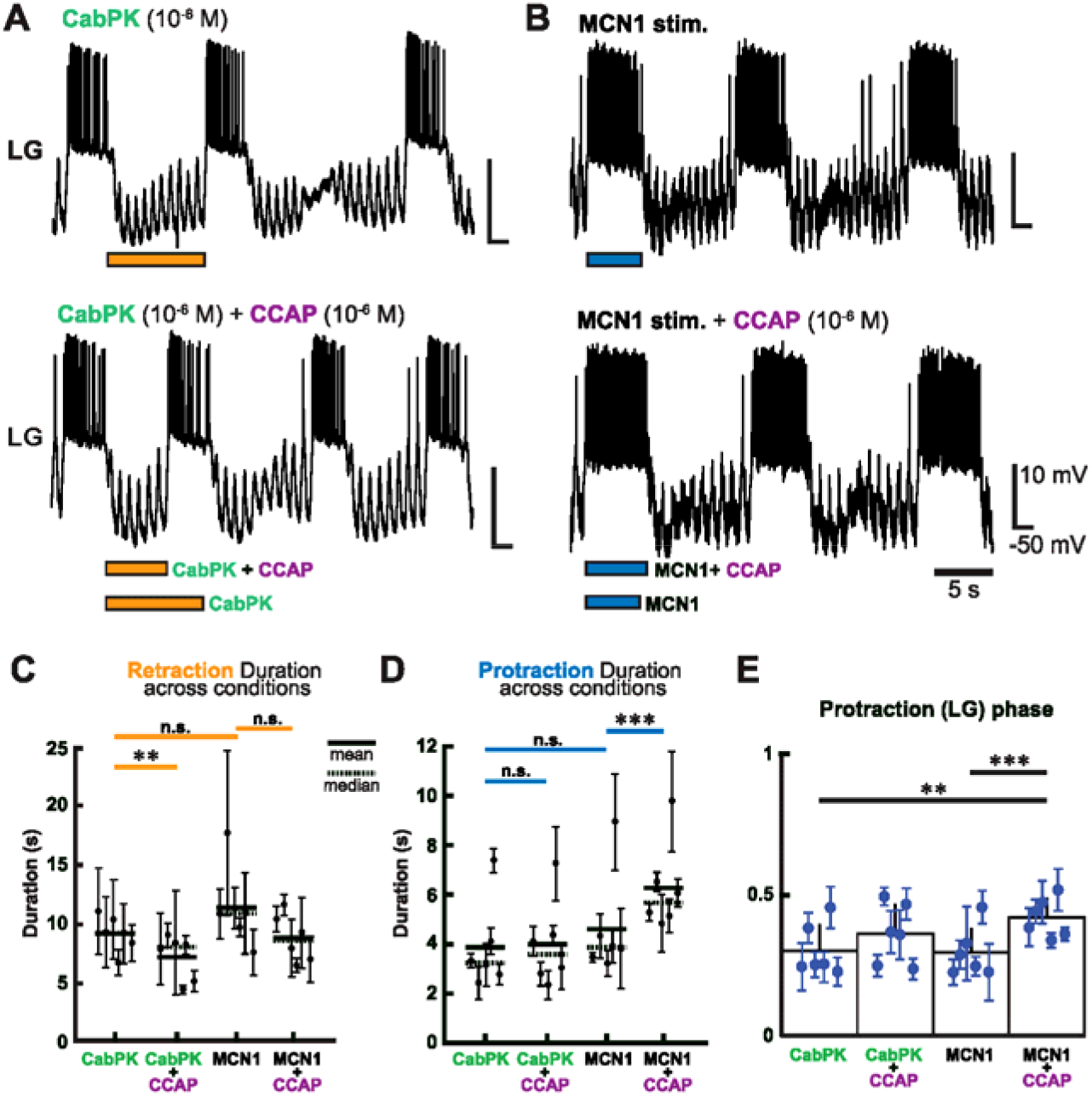
CCAP application (10^−6^ M) selectively decreases the retraction phase duration of the CabPK-gastric mill rhythm and prolongs the protraction duration of the MCN1-gastric mill rhythm. ***A,B*** Example gastric mill rhythms, represented by an intracellular LG recording, driven by (***A***) CabPK application (10^−6^ M) and (***B***) MCN1 stimulation (6 Hz) in the absence (top) and presence (bottom) of CCAP. Orange and blue bars underneath each LG recording denote duration of retraction and protraction phases, respectively. For comparison, orange/blue bars for each control condition are also displayed under the bars for the CCAP modulated rhythms. Values indicated in bottom-right calibration bars pertain to all four traces in (***A***) and (***B***). Recordings in (***A***) and (***B***) are from the same preparation. ***C***, Comparison of mean retraction duration values for each CabPK- and MCN1-gastric mill rhythm without and with CCAP. Retraction duration during CCAP application was different (i.e. decreased) from its control condition only for the CabPK-gastric mill rhythm (repeated paired *t*-tests: CabPK vs CabPK+CCAP, \*\**p* = 0.0088; CabPK vs MCN1, *p* = 0.1534; MCN1 vs MCN1+CCAP, *p* = 0.031; Bonferroni correction: α = 0.0167, n.s. not significantly different, *N* = 6). ***D***, Comparison of mean protraction duration values for both gastric mill rhythms without and with CCAP application. CCAP only altered (prolonged) the MCN1-gastric mill protraction duration (repeated paired *t*-tests: MCN1 vs MCN1+CCAP, \*\*\**p* = 0.00095; CabPK vs CabPK+CCAP, *p* = 0.52; CabPK vs MCN1, *p* = 0.093; Bonferroni correction: α = 0.0167, *N* = 6). For (***C***) and (***D***), we used pair-wise t-tests to compare all conditions except CabPK+CCAP vs MCN1+CCAP, insofar as there was not a biologically informative reason to compare these latter two conditions, and used the Bonferroni correction to account for the accumulated type II error. ***E***, Comparison of mean ± SD LG (i.e. protraction phase) duty cycle across conditions (CabPK: 0.30 ± 0.09, CabPK+CCAP: 0.36 ± 0.10, MCN1: 0.29 ± 0.09, MCN1+CCAP: 0.42 ± 0.07). This duty cycle occupied a larger fraction of the normalized cycle period in the MCN1+CCAP condition than did either CabPK bath application or MCN1 stimulation (MCN1 vs MCN1+CCAP: \*\*\**p* = 0.0005; CabPK vs MCN1+CCAP: \*\**p* = 0.005; all other phase comparisons were not different; repeated paired *t*-tests with a Bonferroni correction: α = 0.0083, *N* = 6).

Despite the selective decrease in gastric mill rhythm retraction duration when CCAP was co-applied with the CabPK, the LG duty cycle was unchanged (**Fig. 2E; Table 3**), presumably due to variability in protraction duration (**Fig. 2D,E**). In contrast, the prolonged protraction phase elicited by CCAP during the MCN1-gastric mill rhythm did increase the LG duty cycle relative to the CabPK- and MCN1(3-6 Hz) conditions in normal saline (**Table 3**). Therefore, while CCAP changes the speed of the CabPK-rhythm, it only consistently altered the LG duty cycle during the MCN1-gastric mill rhythm.

**Table 3.**
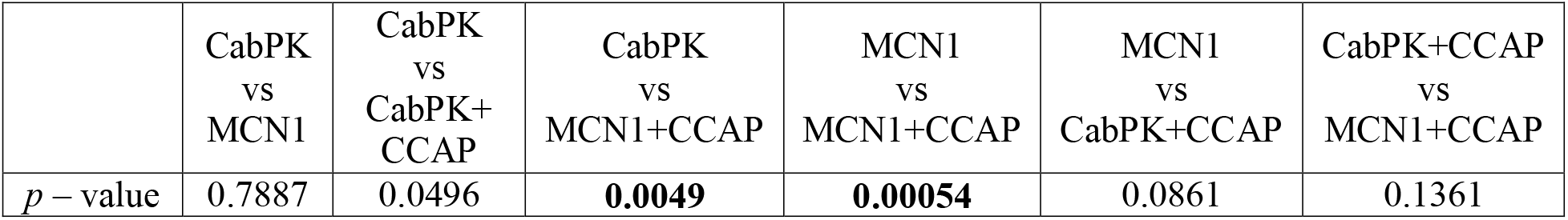
Comparison of LG duty cycle across all conditions. All comparisons performed using repeated paired *t-*tests with a Bonferonni correction α = 0.0083, *N* = 6. Each bolded p-value indicates a significant difference between the two compared conditions.

|  | CabPK<br>vs<br>MCN1 | CabPK<br>vs<br>CabPK+<br>CCAP | CabPK<br>vs<br>MCN1+CCAP | MCN1<br>vs<br>MCN1+CCAP | MCN1<br>vs<br>CabPK+CCAP | CabPK+CCAP<br>vs<br>MCN1+CCAP |
| --- | --- | --- | --- | --- | --- | --- |
| $p$ – value | 0.7887 | 0.0496 | **0.0049** | **0.00054** | 0.0861 | 0.1361 |

### I_MI-CCAP_ is sufficient to selectively decrease CabPK-gastric mill rhythm retraction duration

DeLong et al. (2009b) showed that using the dynamic clamp to inject artificial CCAP-activated I_MI_ (I_MI-CCAP_) into LG was sufficient to mimic the CCAP influence on the MCN1(10-15 Hz)-gastric mill rhythm. More recently, in a computational model of the CabPK-gastric mill rhythm, addition of I_MI-CCAP_ (1-5 pS) into LG also reproduced the biological CCAP reduction in retraction duration during the CabPK-gastric mill rhythm (Rodriguez, 2014). We therefore used the dynamic clamp to determine the likelihood that I_MI-CCAP_ in LG also underlay the distinct CCAP action on the MCN1- and CabPK-gastric mill rhythms.

Dynamic clamp injection of the same artificial I_MI-CCAP_ into LG used by DeLong et al. (2009b) (**Methods**) mimicked many aspects of the CabPK-gastric mill rhythm response to CCAP application (**Fig. 3A**). Specifically, injecting artificial I_MI-CCAP_ caused a comparable decrease (30%) in the CabPK-rhythm retraction duration to that resulting from CCAP application (**Fig. 3B**). Also similar to the CCAP applications, the injected artificial I_MI-CCAP_ (1-3 nA, **Methods**) into LG during CabPK-gastric mill rhythms did not alter protraction duration (**Fig. 3C**).

**Figure 3.**
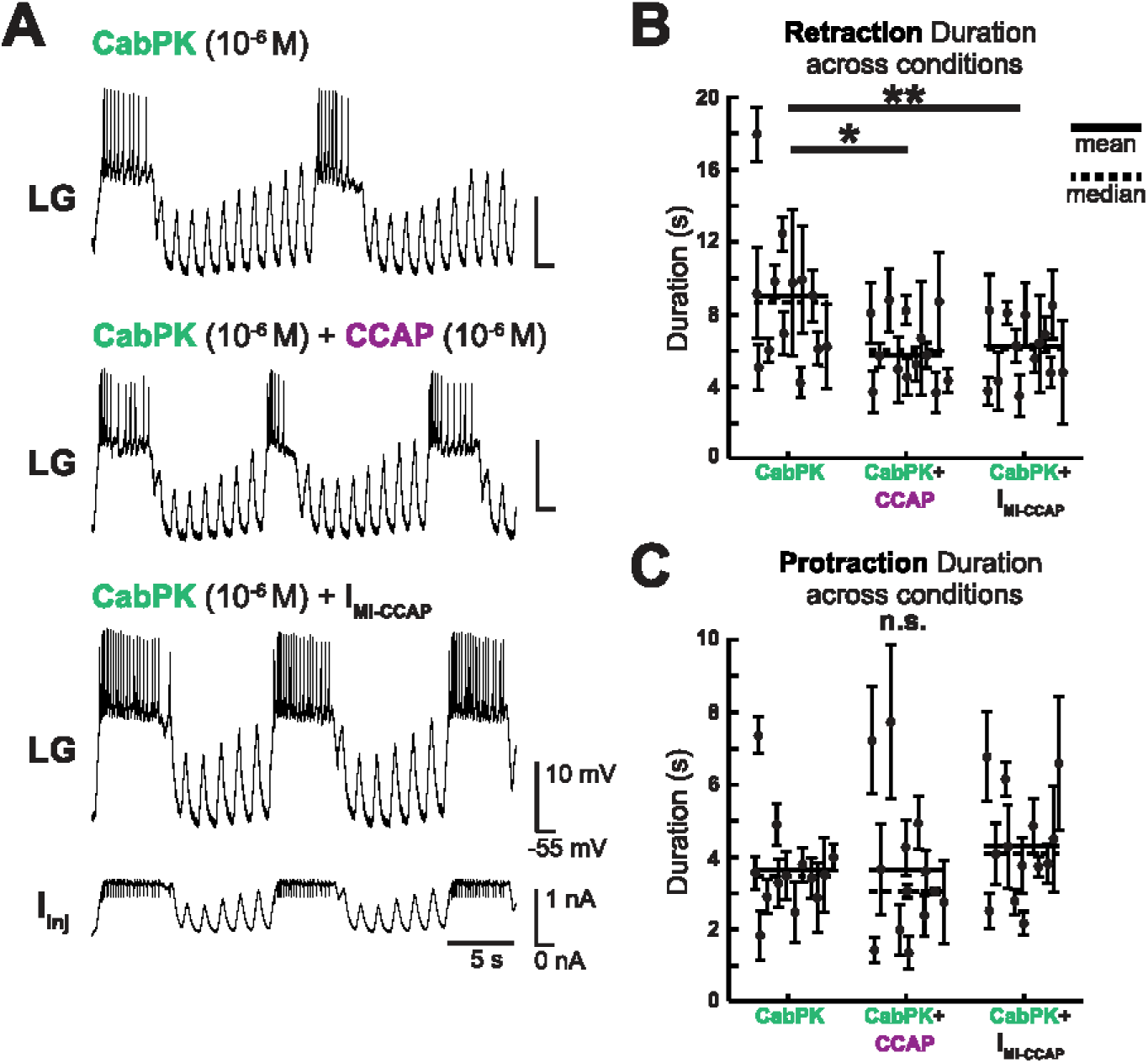
Dynamic clamp-mediated injection of I_MI-CCAP_ into the LG neuron mimics the effect of CCAP bath application on the CabPK-gastric mill rhythm.***A***, Example intracellular recording of the same LG neuron during the CabPK-gastric mill rhythm in CabPK (10^−6^ M) plus saline (top), with co-applied CCAP (10^−6^ M: middle), and with I_MI-CCAP_ injection into LG during CabPK (10^−6^ M) saline application (bottom). The latter recording includes a monitor of the dynamic clamp current injection channel, showing the I_MI-CCAP_ current injected into LG. Calibration bars for LG and time are the same for all conditions. ***B***, Comparison of mean ± SD retraction duration values across experiments for each condition (CabPK: 8.68 ± 3.7 s, CabPK+CCAP: 6.04 ± 1.9 s, CabPK+I_MI-CCAP_: 6.10 ± 1.8 s. RM-ANOVA: df_(within)_ = 12, df_(between)_ = 2, *p* = 0.0001, α = 0.05, *N* = 13; Bonferroni post-hoc: CabPK vs CabPK+CCAP, \**p* = 0.032; CabPK vs CabPK+I_MI-CCAP_, \*\**p* = 0.013; CabPK+CCAP vs CabPK+I_MI-CCAP_, *p* = 0.579, α = 0.05, *N* = 13). ***C***, Comparison of mean ± SD protraction duration values across experiments for each condition (CabPK: 3.65 ± 1.3 s, CabPK+CCAP: 3.64 ± 1.1 s, CabPK+I_MI-CCAP_: 4.31 ± 1.5 s. RM-ANOVA: df_(within)_ = 12, df_(between)_ = 2, (n.s.) *p* = 0.078, α = 0.05, *N* = 13).

Despite the comparable drop in retraction duration, the dynamic clamp injections did not mimic the CCAP-mediated decrease in the CabPK-gastric mill rhythm cycle period (**Table 4**). This was due presumably to a variation in protraction duration resulting from the I_MI-CCAP_ injections. Also consistent with the CCAP applications, these dynamic clamp injections did not alter the number of LG spikes per burst, although the result of these injections did diverge with the effects of CCAP with respect to the LG intraburst spike frequency (RM-ANOVA, df_(within)_ = 12, df_(between)_ = 2: intraburst spike frequency, *p* = 0.87; spikes per burst, *p* = 0.079, α = 0.05; *N* = 13) (**Table 4**).

**Table 4.** Comparison of CabPK-gastric mill rhythm parameters under different conditions during dynamic clamp experiments. PK = CabPK II; I_MI_ = I_MI-CCAP_; −I_MI_ = −I_MI-CCAP_; Column definitions as in Table 1. All values are mean ± SD, *N* = 13. Statistical analysis, Rows 1-3 (associated with **Fig. 3**): Cycle period: RM-ANOVA: df_(within)_ = 12, df_(between)_ = 2, *p* = 0.006; Bonferroni post-hoc test; CabPK vs CabPK+CCAP: \**p* = 0.031; CabPK vs CabPK+I_MI-CCAP_: *p* = 0.065; CabPK+CCAP vs CabPK+I_MI-CCAP_: *p* = 0.796; α = 0.05, *N* = 13. Number of LG spikes per burst: RM-ANOVA, *p* = 0.08; LG spike frequency, *p* = 0.87, α = 0.05, *N* = 13. Rows 1-2, 4 (assoc. w/**Fig. 4**): RM-ANOVA, df_(within)_ = 12, df_(between)_ = 2, *p* = 0.006; Bonferroni post-hoc test; CabPK vs CabPK+CCAP: * *p* = 0.031; CabPK vs CabPK+CCAP+ −I_MI-CCAP_: *p* = 0.685; CabPK+CCAP vs CabPK+CCAP+ −I_MI-CCAP_: ** *p* = 0.012; α = 0.05, *N* = 13. RM-ANOVA for number of LG spikes per burst, *p* = 0.25; LG spike frequency, *p* = 0.38, α = 0.05, *N* = 13. See Figures 3 and 4 for the statistical comparisons of the data in columns 1 and 2.

|  | Prot. (s) | Ret. (s) | Cycle Period (s) | LG<br>#Spikes/Burst | LG Spike Freq.<br>(Hz) |
| --- | --- | --- | --- | --- | --- |
| PK | 3.65 ± 1.3 | 8.68 ± 3.6 | 12.33 ± 4.3 | 19.80 ± 7.4 | 5.94 ± 2.7 |
| PK+CCAP | 3.64 ± 2.0 | 6.05 ± 1.9 | 9.69 ± 3.5 | 21.90 ± 14.0 | 6.25 ± 2.2 |
| PK+I<sub>MI</sub> | 4.31 ± 1.5 | 6.10 ± 1.8 | 10.42 ± 2.8 | 27.04 ± 12.2 | 6.14 ± 1.5 |
| PK+CCAP+ -I<sub>MI</sub> | 3.70 ± 2.2 | 8.30 ± 3.2 | 12.00 ± 4.4 | 25.00 ± 17.4 | 6.77 ± 2.1 |

### Injecting negative I_MI-CCAP_ into LG nullifies the CCAP modulation of the CabPK-gastric mill rhythm

Given the success of injecting artificial I_MI-CCAP_ into LG to mimic the influence of CCAP application on the CabPK-gastric mill rhythm, we next injected into LG a negative conductance version of g_MI-CCAP_ (−g_MI-CCAP_) during CCAP application, to determine if it would nullify the CCAP influence on the CabPK-gastric mill rhythm. **Figure 4A** shows that this approach did successfully occlude the CCAP effect. For example, the addition of artificial −g_MI-CCAP_ did nullify the CCAP reduction in retraction duration **(Fig. 4B)**. Consequently, the injected −g_MI-CCAP_ also nullified the CCAP reduction in cycle period (**Table 4**).

**Figure 4.**
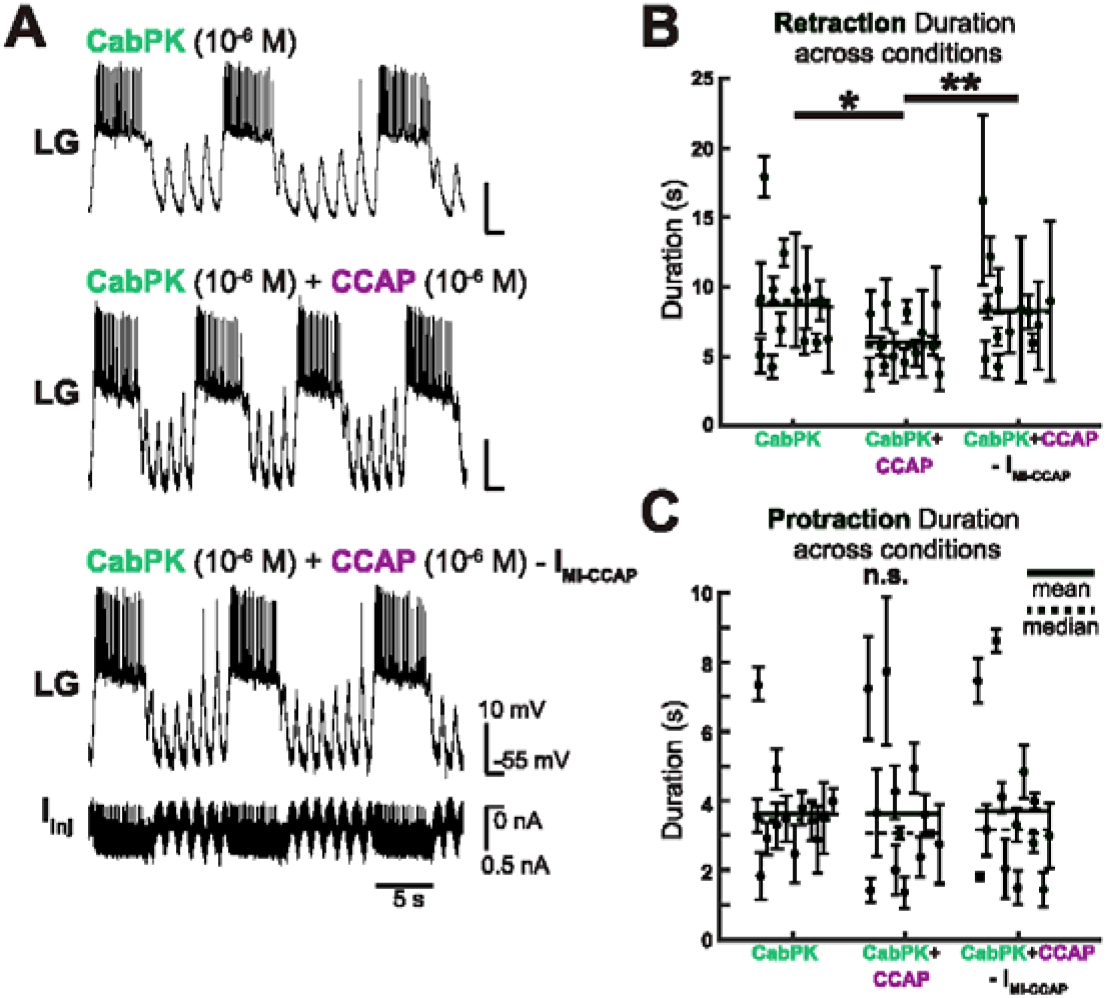
Dynamic clamp injection of the negative version of I_MI-CCAP_ into the LG neuron nullifies the effect of bath applied CCAP on the CabPK-gastric mill rhythm. ***A***, Example intracellular recording of the same LG neuron during the CabPK-gastric mill rhythm in CabPK (10^−6^ M) saline (top), with applied CCAP (10^−6^ M) (middle), and with applied CCAP plus injection of −I_MI-CCAP_ (bottom). The latter condition includes a monitor of the current injection channel showing the −I_MI-CCAP_ injection into LG. Calibration bars for LG and time are the same for all panels. ***B***, Comparison of mean ± SD retraction duration during each condition (CabPK: 8.68 ± 3.7 s, CabPK+CCAP: 6.05 ± 1.9 s, CabPK+CCAP+ −I_MI-CCAP_: 8.30 ± 3.2 s; RM-ANOVA: df_(within)_ = 12, df_(between)_ = 2, *p* = 0.003, α = 0.05, *N* = 13; Bonferroni post-hoc: CabPK vs CabPK+CCAP, \**p* = 0.032; CabPK vs CabPK+ CCAP+ −I_MI-CCAP_, *p* = 0.613; CabPK+CCAP vs CabPK+CCAP+ −I_MI-CCAP_, \*\**p* = 0.014, α = 0.05, *N* = 13). ***C***, Comparison of mean ± SD protraction duration values across experiments for each condition (CabPK: 3.65 ± 1.3 s, CabPK+CCAP: 3.64 ± 1.1 s, CabPK+CCAP+ −I_MI-CCAP_: 3.70 ± 2.2 s. RM-ANOVA: df_(within)_ = 12, df_(between)_ = 2, *p* = 0.981, α = 0.05, *N* = 13).

Furthermore, injecting artificial −g_MI-CCAP_ did not artifactually alter the CabPK-rhythm parameters that were not altered by CCAP application, such as protraction duration (**Fig. 4C**) and the number of LG spikes per burst, but it did nullify the CCAP-mediated increase in the LG intraburst spike frequency (RM-ANOVA: df_(within)_ = 12, df_(between)_ = 2; intraburst spike frequency, *p* = 0.376, spikes per burst, *p* = 0.251; α = 0.05, *N* = 13) (**Table 4**).

### The pyloric rhythm cycle frequency is equivalent during CabPK and CabPK+CCAP applications

The pyloric rhythm pacemaker neuron AB sets the timing of the MCN1- and CabPK-gastric mill rhythms, with AB activity regulating the MCN1-rhythm and being necessary for the CabPK-rhythm (**Fig. 1B**) (Nadim et al., 1998; Bartos et al., 1999; Saideman et al., 2007a). This AB neuron role results from each LG neuron burst initiating during an episode of AB inhibition of Int1. Thus, during these gastric mill rhythms each protraction phase onset (i.e. LG burst onset) is coupled to the start of a pyloric cycle (i.e. AB burst) (Nadim et al., 1998; Bartos et al., 1999; Saideman et al., 2007a).

In the isolated STG, in the absence of MCN1 stimulation or CabPK application, CCAP elicits a cycle frequency-dependent increase in the pyloric rhythm, increasing the rhythm speed when the control rhythm is ≤0.5 Hz (Weimann et al., 1997). However, the CCAP influence on the MCN1(10-15 Hz)-gastric mill rhythm was not an indirect consequence of its actions on the pyloric rhythm (Kirby and Nusbaum, 2007). Given the necessary role of AB on CabPK rhythm generation, we determined whether CCAP application altered the pyloric rhythm during the CabPK-gastric mill rhythm and, if so, whether this change influenced the reduction in retractor phase duration of the CabPK-rhythm with CCAP co-application. We found that the pyloric cycle frequency was not altered by CCAP during the CabPK-gastric mill rhythm (CabPK: 0.72 ± 0.34 Hz, CabPK + CCAP: 0.84 ± 0.29 Hz; paired *t*-test: *p* =0.72, α = 0.05, *N* = 9).

We anticipated that the number of pyloric cycles per gastric mill cycle would be decreased when CCAP was applied during the CabPK-gastric mill rhythm. This expectation was based on these two rhythms being coordinated during the CabPK-gastric mill rhythm, the pyloric cycle frequency being the same during CabPK-gastric mill protraction and retraction and not changing when CCAP was applied, and the gastric mill cycle period being reduced (Saideman et al., 2007a; this paper). This expectation was indeed realized (# pyloric cycles/gastric mill rhythm cycle during: CabPK, 11.59 ± 6.0; CabPK+CCAP: 9.28 ± 4.4; paired *t*-test: *p* = 0.013, α = 0.05, *N* = 9). Thus, the added depolarizing drive of LG due to CCAP allows each LG burst to initiate a few pyloric cycles sooner than during CabPK alone.

### GPR regulation of the MCN1-gastric mill rhythm is MCN1 firing rate-dependent

In the isolated *C. borealis* STG, stimulating the muscle stretch-sensitive GPR neurons evokes both excitatory and inhibitory responses from gastric mill and pyloric circuit neurons (Katz et al., 1989; Katz and Harris-Warrick, 1989, 1990; Beenhakker et al., 2005). With respect to the gastric mill circuit, GPR provides ionotropic excitation to Int1, metabotropic inhibition to the LG neuron and MCN1_STG_, and metabotropic excitation to DG (**Fig. 1D**) (Kiehn and Harris-Warrick, 1992; Beenhakker et al., 2005; DeLong et al., 2009a). GPR is activated by stretch of the protractor muscles innervated by its dendrites, and thus is active during the gastric mill retraction phase (Katz et al., 1989; Birmingham et al., 1999). In some preparations, GPR also generated rhythmic bursts that were independent of muscle stretch (Birmingham et al., 1999). During this latter condition, GPR activity would occur during either gastric mill retraction or protraction.

GPR stimulation during the MCN1(10-15 Hz)-gastric mill rhythm selectively prolongs retraction, having no effect on protraction duration whether it was stimulated during protraction or retraction (DeLong et al., 2009a). Similarly, GPR stimulation (6-8 Hz) prolonged the MCN1(3-6 Hz)-gastric mill rhythm retraction phase by 37% (**Fig. 5A,B**). This GPR stimulation did not alter the protraction duration of the subsequent LG burst (RM-ANOVA: df_(within)_ = 5, df_(between)_ = 2, *p* = 0.271, α = 0.05, *N* = 6) (**Table 5**).

**Figure 5.**
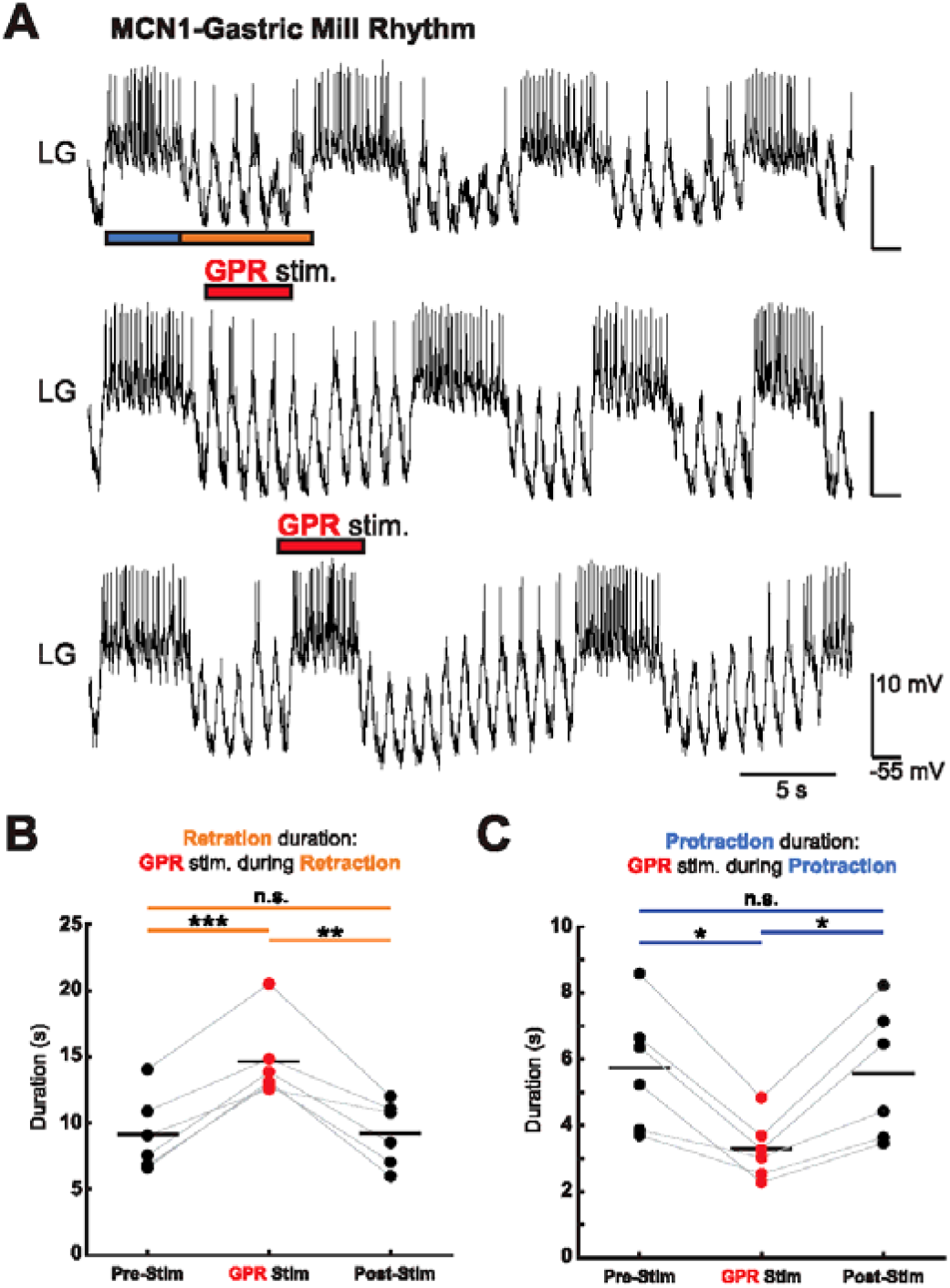
GPR stimulation (6-8 Hz) during the MCN1(3-6 Hz)-gastric mill rhythm protraction or retraction phase selectively prolongs retraction and shortens protraction, respectively. ***A***, Example intracellular recording of the same LG neuron during the MCN1-gastric mill rhythm without GPR stimulation (top) and with GPR stimulation during one retraction phase (middle) or protraction phase (bottom). Blue and orange bars indicate protraction and retraction phase, respectively. Note that GPR stimulation (6 Hz) during retraction prolonged that phase (middle trace) while stimulating it during protraction shortened that phase (bottom). Red bars indicate timing and duration of GPR stimulation. All traces in (***A***) are from the same preparation and occurred within an 8-minute window. Values for LG and time calibration bars are the same for all recordings. ***B***, Comparison of the MCN1-gastric mill rhythm retraction duration (mean ± SD) before (Pre-Stim.: 9.17 ± 2.9 s), during (GPR Stim.: 14.64 ± 3.0 s), and after (Post-Stim.: 9.24 ± 2.4 s) GPR stimulation during retraction (RM-ANOVA: df_(within)_ = 5, df_(between)_ = 2,*p* < 0.001, Bonferroni post-hoc: Pre-vs During GPR stim., \*\*\**p* = 0.001; Pre-vs Post-GPR stim., *p* = 1; GPR stim. vs Post-GPR stim., \*\**p* = 0.008; α = 0.05, *N* = 6). Lines between conditions link mean values from the same experiment. ***C***, Comparison of the mean ± SD protraction duration across experiments for the MCN1-gastric mill rhythm before (Pre-Stim: 5.73 ± 1.9 s), during (GPR Stim.: 3.27 ± 0.9 s), and after (Post-Stim.: 5.56 ± 2.0 s) GPR stimulation during protraction (RM-ANOVA: df_(within)_ = 5, df_(between)_ = 2, *p* < 0.001, Bonferroni post-hoc: Pre-vs During GPR stim., \**p* = 0.011; Pre-vs Post-GPR stim., *p* = 1; GPR stim. vs Post-GPR stim., \**p* = 0.015; α = 0.05, *N* = 6). Lines between conditions link mean values from the same experiment. Black horizontal lines indicate the means for each condition in (***B***) and (***C***).

**Table 5.**
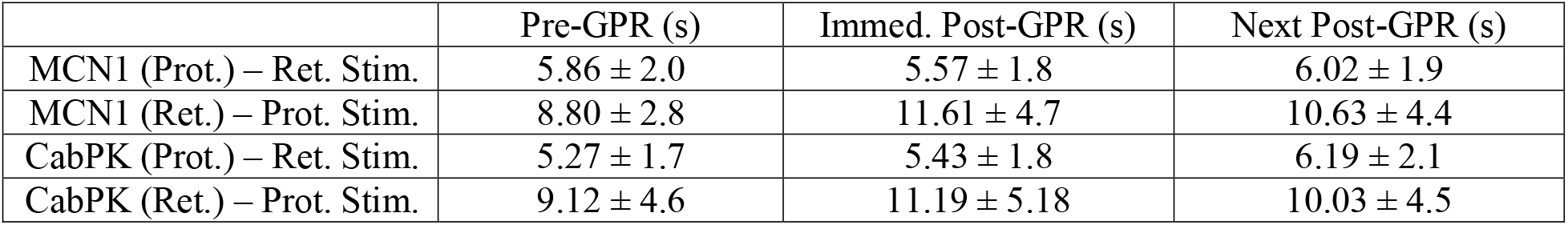
Mean MCN1- and CabPK-gastric mill phase durations bracketing the phase during which GPR was stimulated. Pre-GPR, phase duration immediately before GPR stimulation; Immed. Post-GPR, phase duration immediately after GPR stimulation; Next Post-GPR, phase duration after Immed. Post-GPR. In each case, values (mean ± SD; MCN1-rhythm *N* = 6, CabPK-rhythm: *N* = 16) come from the phase of the cycle not stimulated (e.g. for the first row, values are from protraction duration when GPR was stimulated during retraction).

|  | Pre-GPR (s) | Immed. Post-GPR (s) | Next Post-GPR (s) |
| --- | --- | --- | --- |
| MCN1 (Prot.) – Ret. Stim. | 5.86 ± 2.0 | 5.57 ± 1.8 | 6.02 ± 1.9 |
| MCN1 (Ret.) – Prot. Stim. | 8.80 ± 2.8 | 11.61 ± 4.7 | 10.63 ± 4.4 |
| CabPK (Prot.) – Ret. Stim. | 5.27 ± 1.7 | 5.43 ± 1.8 | 6.19 ± 2.1 |
| CabPK (Ret.) – Prot. Stim. | 9.12 ± 4.6 | 11.19 ± 5.18 | 10.03 ± 4.5 |

However, stimulating GPR (6-8 Hz) during the MCN1(3-6 Hz)-rhythm protraction phase reduced protraction duration by 43% (**Fig. 5A,C**). These stimulations during protraction did not affect the subsequent retraction phase duration (RM-ANOVA: df(_within_) = 5, df_(between)_ = 2, α = 0.05, *p* = 0.022; Bonferroni post-hoc: Pre-GPR vs Immed. Post-GPR, *p* = 0.091; Pre-GPR vs Next Post-GPR, *p* = 0.35; Immed. Post-GPR vs Next Post-GPR, *p* = 0.44; α = 0.05, *N* = 6) (**Table 5**).

The observed difference between stimulating GPR (6-8 Hz) during the protraction phase of the MCN1(10-15 Hz)-rhythm (DeLong et al., 2009a) and the MCN1(3-6 Hz)-rhythm (this paper) is likely due to do a greater I_MI_ build-up in LG resulting from higher MCN1 stimulation frequencies. Increased I_MI_ increases LG excitability and likely occludes the inhibitory drive from GPR. For example, the LG intraburst spike frequency is higher during 10-15 Hz stimulation (8.29 ± 3.3 Hz) than 3-6 Hz stimulation (4.52 ± 0.8 Hz) (Independent sample *t*-test of unequal variance: *p* = 0.009; *N* = 5 & 6 respectively).

GPR also excites the DG neuron, which in turn inhibits LG (**Fig. 1D**). This connectivity suggested that the GPR-mediated increase in retraction duration during the MCN1-gastric mill rhythm might result from enhanced DG activity preventing LG from escaping Int1 inhibition. DeLong et al. (2009a) showed that this was not the case, as hyperpolarizing DG did not alter the effect of GPR stimulation during retraction. Therefore, DG was hyperpolarized for the data collected in **Figure 5**. However, to assess whether GPR stimulation during the MCN1(3-6 Hz)-rhythm was also influenced by DG activity, we performed additional experiments to determine the effect of GPR (6-8 Hz) stimulation on this gastric mill rhythm when DG was active.

When GPR was stimulated during retraction with DG active, the same qualitative effects were observed as when DG was silenced by hyperpolarization. For example, retraction duration was prolonged (55%) during the GPR stimulation relative to the preceding and following cycles (**Table 6**). This effect was similar to that occurring with DG hyperpolarized [Retraction duration difference values (GPR stim. minus pre-GPR stim.): DG_Active_: 7.84 ± 2.8 s, *N* = 4; DG_Inactive_: 5.47 ± 1.4 s, *N* = 6; Two-sample *t*-test: *p* = 0.11]. This was also the case with respect to the lack of change in the subsequent protraction duration (Pre-GPR: 5.43 ± 0.7 s, Immed. Post-GPR: 5.49 ± 0.6 s, Next Post-GPR: 5.75 ± 0.9 s. RM-ANOVA: df_(within)_ = 3, df_(between)_ = 2, *p* = 0.31 α = 0.05, *N* = 4).

**Table 6.** Mean gastric mill phase durations during the MCN1-rhythm phase in which GPR was stimulated with DG hyperpolarized. Values (mean ± SD) in the second row are retraction durations before (Pre-GPR), during (GPR Stim.), and after (Post-GPR) the retraction phase where GPR was stimulated (***N*** = 4). Values in the third row correspond with protraction durations when GPR was stimulated during protraction (***N*** = 4). Comparison of retraction durations: RM-ANOVA: df_(within)_ = 3, df_(between)_ = 2, *p* = 0.001; Bonferroni post-hoc: Pre-GPR vs GPR stim., * *p* = 0.035; GPR stim. vs Post-GPR; ** *p* = 0.009; Pre-GPR vs Post-GPR, *p* = 1; α = 0.05, *N* = 4. Comparison of protractor durations: RM-ANOVA: df_(within)_ = 3, df_(between)_ = 2, α = 0.05, *p* = 0.001; Bonferroni post-hoc: Pre-GPR stim. vs GPR stim., * *p* = 0.044; GPR stim. vs Post-GPR stim.; * *p* = 0.041; Pre-GPR stim. vs Post-GPR stim., *p* = 1; α = 0.05, *N* = 4.

|  | Pre-GPR (s) | GPR Stim. (s) | Post-GPR (s) |
| --- | --- | --- | --- |
| MCN1 – Ret. Stim | 6.35 ± 2.7 | 14.2 ± 4.5 | 7.07 ± 2.9 |
| MCN1 – Prot. Stim | 5.43 ± 0.7 | 2.5 ± 1.8 | 5.42 ± 0.7 |

It was also possible that the GPR truncation of protraction duration during the MCN1(3-6 Hz)-rhythm when GPR stimulation occurred during protraction resulted from it prematurely activating DG, enabling DG to prematurely inhibit LG. With DG active, GPR stimulation during protraction shortened the LG burst by 54% (**Table 6**). This result, however, was comparable to that resulting from the same GPR stimulation with DG hyperpolarized (Protraction duration difference values: DG_Active_: 2.93 ± 1.2 s, *N* = 4; DG_Inactive_: 2.46 ± 1.2 s, *N* = 6; Two-sample *t*-test: *p* = 0.55).

### GPR has comparable effects on the MCN1- and CabPK-gastric mill rhythm

Stimulating GPR (6-8 Hz) during the CabPK-gastric mill rhythm, without suppressing DG neuron activity, had similar effects to those during the MCN1(3-6 Hz)- and MCN1(10-15 Hz)-gastric mill rhythm (DeLong et al., 2009b; this paper). Stimulating GPR during the CabPK-rhythm retraction phase consistently prolonged this phase, by ~45% (**Fig. 6A,B**). Also consistent with stimulating GPR during the MCN1-gastric mill retraction phase, GPR stimulation did not alter the subsequent protraction duration (RM-ANOVA: *p* = 0.093, α = 0.05, *N* = 16) (**Table 5**).

**Figure 6.**
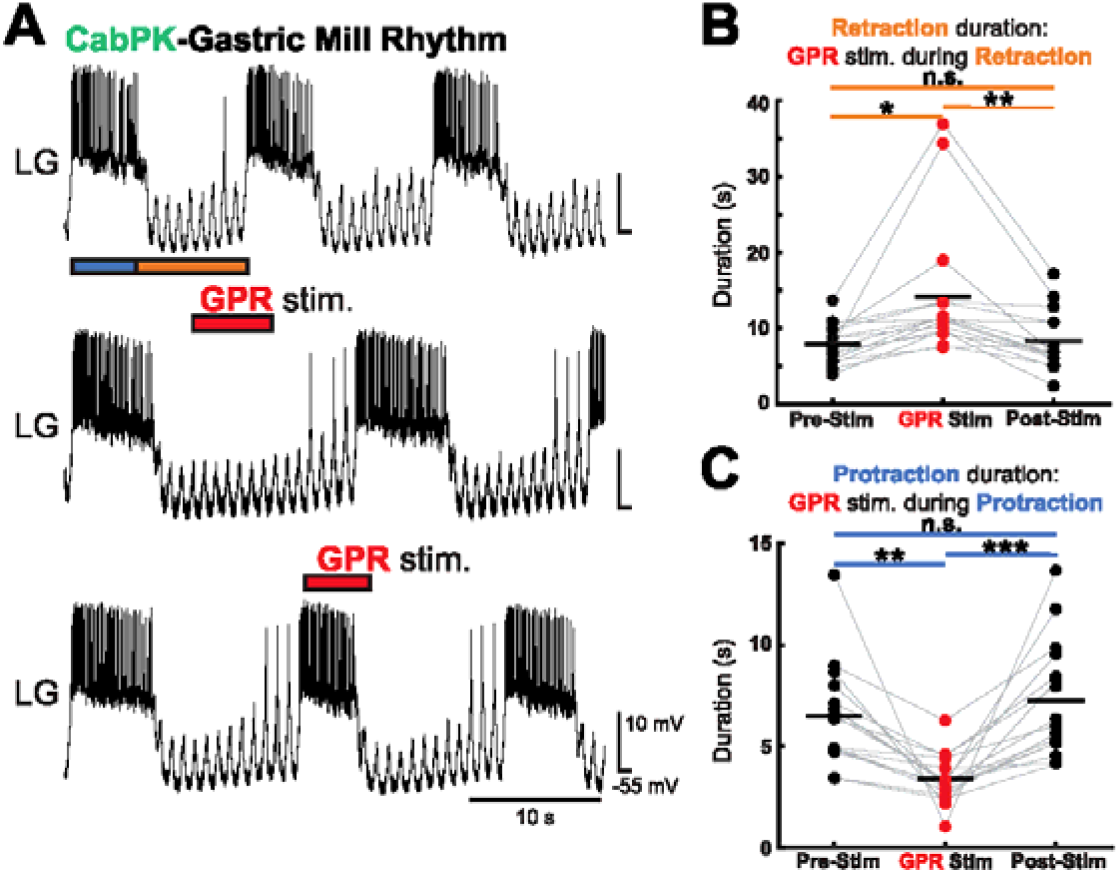
GPR stimulation (6-8 Hz) during the CabPK-gastric mill rhythm protraction or retraction phase selectively prolongs retraction and shortens protraction, respectively. ***A***, Example intracellular recordings of the same LG neuron during the CabPK (10^−6^ M)-gastric mill rhythm without GPR stimulation (top) and with GPR stimulation (6 Hz, red bars) during retraction (middle) or protraction (bottom). Blue and orange bars indicate protraction and retraction phases, respectively. Note that GPR stimulation selectively prolonged retraction when stimulated in that phase, and it selectively shortened protraction duration when stimulated during that phase. All traces in (***A***) are from the same preparation and occurred within an 8-minute window. ***B***, Comparison of the mean ± SD retraction duration of the CabPK-gastric mill rhythm across experiments before (Pre-Stim.: 7.89 ± 2.6 s), during (GPR Stim.: 14.2 ± 8.8 s), and after (Post-Stim.: 8.3 ± 3.9 s) GPR stimulation during retraction (RM-ANOVA: df_(within)_ = 15, df_(between)_ = 2, *p* < 0.001, Bonferroni post-hoc: Pre-vs During GPR stim., \**p* = 0.015; Pre-vs Post-GPR stim.,*p* = 1; GPR stim. vs Post-GPR stim., \*\**p* = 0.006; α = 0.05; *N* = 16). Lines between conditions link mean values from the same experiment. ***C***, Comparison of the mean ± SD protraction duration of the CabPK-gastric mill rhythm across experiments before (Pre-Stim.: 6.49 ± 2.5 s), during (GPR Stim.: 3.36 ± 1.2 s), and after (Post-Stim.: 7.25 ± 2.8 s) GPR stimulation during protraction (RM-ANOVA: df(_within_) = 15, df_(between)_ = 2, *p* < 0.001, Bonferroni post-hoc: Pre-vs During GPR stim., \*\**p* = 0.002; Pre-vs Post-GPR stim., *p* = 0.14; GPR stim. vs Post-GPR stim., \*\*\**p* = 0.001; α = 0.05; *N* = 16). Lines between conditions link mean values from the same experiment. Black horizontal lines indicate the means for each condition in (***B***) and (***C***).

Similar to the MCN1(3-6 Hz)-rhythm, when GPR was stimulated during the protraction phase of the CabPK-gastric mill rhythm, protraction duration was shortened by 48%, and the GPR effect was limited to the protraction phase in which it was stimulated (**Fig. 6A,C**) (**Table 5**). We also examined this GPR influence with DG hyperpolarized, as during the MCN1-rhythm. Under this condition, GPR stimulation during protraction still reduced protraction duration (paired *t*-test: *p* = 0.021, α = 0.05, *N* = 5; data not shown).

We wanted to rule out the possibility that the effect of GPR stimulation during the CabPK-rhythm resulted from back-activation of MCN1_STG_ by CabPK application and therefore a consequence of the GPR inhibition of MCN1_STG_ (DeLong et al., 2009a). CabPK (10^−6^ M) application elicits a transient and often weak activation of MCN1 in ~10% of preparations (Saideman et al., 2007b). In 9 of 16 experiments, we had an intracellular recording of LG and determined the frequency of electrical EPSPs (eEPSPs) from MCN1 to LG during CabPK bath application (1.43 ± 0.8 Hz; Range: 0-2.3 Hz). The effectiveness of GPR stimulation during the CabPK-rhythm retraction phase was comparable when MCN1 was silent or weakly active (≤ 1 Hz; 3/9 experiments), where GPR prolonged retraction by 31%, and when MCN1 was more active (1-2.3 Hz; 6/9 exps), during which GPR prolonged retraction by 29%. These MCN1 firing frequencies are well below the threshold for stimulating a MCN1-gastric mill rhythm (Kirby and Nusbaum, 2007). Thus, the effect of retraction phase GPR stimulation during the CabPK-rhythm likely occurs via a distinct synaptic pathway from that occurring during the MCN1-rhythm.

To determine whether GPR stimulation during the CabPK-gastric mill rhythm retraction phase prolonged this phase by indirectly inhibiting LG, via its excitation of DG, we performed additional GPR stimulations with DG hyperpolarized. Because the variance in retraction duration during the GPR stimulations was large for this data set (Bartlett’s test, *p* = 0.0045, *N* = 6), a nonparametric test was used to compare retractor duration before and during GPR stimulation with DG hyperpolarized (Wilcoxon Signed-Rank test, *p* = 0.031, α = 0.05, *N* = 6; data not shown). This result indicated that the DG inhibition of LG was not necessary for enabling GPR to prolong the retraction duration during the CabPK-gastric mill rhythm. This manipulation also did not affect the subsequent protraction duration (Wilcoxon Signed-Rank test, *p* = 1, α = 0.05, *N* = 6; data not shown).

### GPR likely has a direct inhibitory synapse on LG

During the CabPK-gastric mill rhythm, GPR stimulation during protraction could have limited protraction duration by inhibiting LG and/or prematurely activating Int1 (**Fig. 1D**). GPR does directly excite Int1, but whether its inhibition of LG is direct was not known (Beenhakker et al., 2005; DeLong et al., 2009a). Insofar as GPR excites Int1 and Int1 inhibits LG, the GPR inhibition of LG might have been entirely indirect. Int1 makes a glutamatergic inhibitory synapse onto LG, so we examined the GPR influence on LG with glutamate receptor-mediated inhibition in the STG blocked with bath application of picrotoxin (PTX: 10^−5^ M; **Methods**) (Marder and Paupardin-Tritsch, 1978; Bidaut, 1980; Marder and Eisen, 1984).

In the presence of PTX, we injected constant amplitude and duration depolarizing current steps into LG, each of which elicited tonic firing (**Fig. 7A**). During these rhythmic depolarizations, GPR stimulation hyperpolarized LG by ~2.7 mV (**Fig. 7A,B**). Moreover, during these GPR stimulations LG exhibited 51% fewer spikes per depolarization (**Fig. 7C**). This drop in the number of spikes per depolarization was accompanied by a 61% reduction in LG spike frequency (Pre-GPR: 8.48 ± 3.9 Hz, GPR: 3.27 ± 1.2 Hz, Post-GPR stim.: 7.43 ± 2.9 Hz; repeated paired *t*-tests with a Bonferroni correction: α = 0.0167, Pre-GPR vs GPR, *p* = 0.0079; GPR vs Post-GPR: *p* = 0.0041; Pre-GPR vs Post-GPR, *p* = 0.07, *N* = 7). These GPR stimulations elicited no unitary IPSPs in LG, and the resulting inhibitory response outlasted the stimulation by several seconds, suggesting that this GPR action was metabotropic. For example, the first depolarizing current injection into LG after GPR stimulation still elicited fewer spikes (Pre-GPR: 10.5 ± 3.6, Immed. Post-Stim.: 7.08 ± 2.6; paired *t*-test: *p* = 0.0012, *N* = 7) at a reduced firing frequency (Pre-GPR: 8.49 ± 3.9 Hz, Immed. Post-Stim.: 4.77 ± 1.6 Hz; paired *t*-test: *p* = 0.036, *N* = 7) relative to pre-GPR stimulation. These results support a direct inhibitory synapse from GPR to LG, which could underlie the LG response to GPR stimulation during both the MCN1(3-6 Hz)- and the CabPK-gastric mill rhythm.

**Figure 7.**
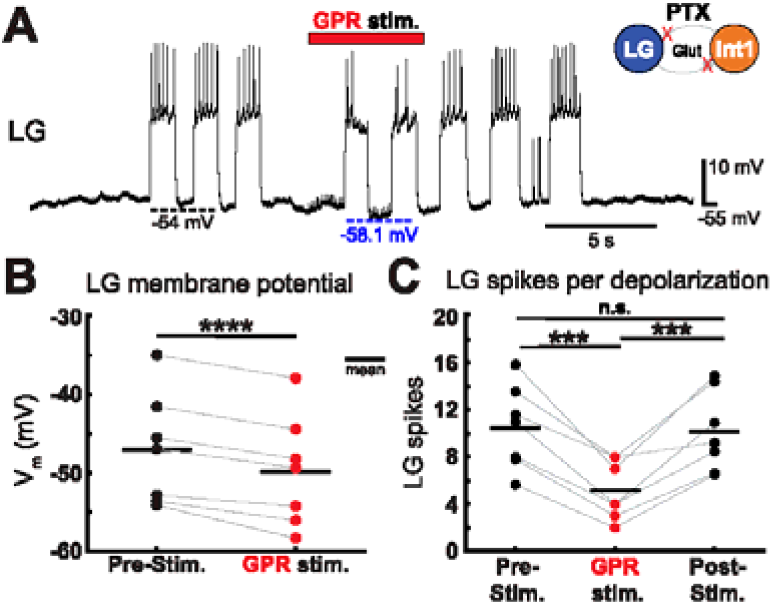
GPR stimulation (6-8 Hz) continues to inhibit the LG neuron when LG is functionally isolated from STG inputs with picrotoxin (PTX: 10^−5^ M) saline. PTX suppresses ionotropic glutamatergic inhibition in the STG, including the reciprocal inhibitory synapses between LG and Int1 (inset). Using two-electrode current clamp, suprathreshold depolarizing current steps of constant amplitude and duration were repeatedly injected into LG to determine the LG response to GPR stimulation. ***A***, Example intracellular recording of the LG neuron while current steps are injected and GPR is stimulated (red bar). Stimulation artifacts are evident in the LG recording. Note that GPR-mediated weakening of LG activity outlasts the GPR stimulus duration and that LG membrane potential hyperpolarization is delayed compared to the onset of GPR stimulation, and then repolarizes gradually after GPR stimulation is terminated. Dashed lines indicate baseline V_m_ in LG before (black) and during (blue) GPR stimulation. ***B***, Comparison of the mean ± SD baseline LG membrane potential (V_m_) before (black: −47.1 ± 7.1 mV) and during (red: −49.8 ± 7.1 mV) GPR stimulation (paired t-test: \*\*\*\**p* = 0.0001, α = 0.05; *N* = 7), as represented by the black and blue dashed lines in (***A***). Lines connect values from the same experiment. ***C***, The mean ± SD number of LG spikes per depolarization per experiment before (Pre-Stim.: 10.5 ± 3.6), during (GPR Stim.: 5.14 ± 2.5), and after (Post-Stim.: 10.1 ± 3.4) GPR stimulation (repeated paired t-tests: Pre-Stim vs GPR: \*\*\**p* = 0.0004, GPR vs Post-Stim.: \*\*\**p* = 0.001, Pre-vs Post-Stim.: *p* = 0.49; α = 0.0167 (Bonferroni correction); *N* = 7). Lines connect values from the same experiment.

## Discussion

### Comparable rhythms do not necessarily respond comparably to the same input

We examined whether providing two degenerate circuit states with the same input would expose their distinct underlying states by eliciting different output patterns. To this end, we used a peptide hormone (CCAP) and a sensory feedback neuron (GPR) to challenge two different gastric mill circuits in the crab STG that generate comparable gastric mill rhythms via distinct cellular and synaptic mechanisms (Saideman et al., 2007a; Rodriguez et al., 2013). The distinct underlying state of these two circuits was indeed exposed as they generated divergent responses to CCAP application (**Fig. 8A**). However, they generated equivalent responses to GPR stimulation, despite the fact that this similar response occurred by GPR using a different synapse to influence each circuit (**Fig. 8B**). Therefore, using responses to exogenous input as a litmus test for network degeneracy can be problematic without direct access to the relevant circuit(s).

**Figure 8.**
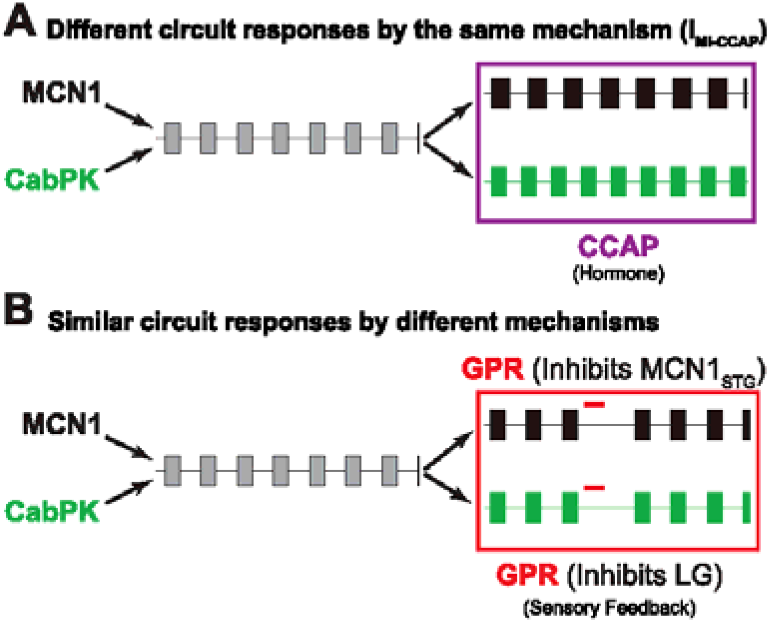
Schematic illustrating that degenerate circuits can respond differently or similarly to a shared input. ***A***, Application of the peptide hormone CCAP altered the MCN1-gastric mill rhythm by selectively prolonging the protraction phase (black pattern), while it increased the speed of the CabPK-gastric mill rhythm by selectively reducing retraction duration (green pattern). ***B***, Stimulating the sensory neuron GPR during the gastric mill retraction phase slowed both gastric mill rhythms by selectively prolonging retraction, albeit via different synapses.

Similar network output that arises from distinct circuit states can diverge in surprising ways in response to a modulatory influence (Grashow et al., 2009, 2010; Marder et al., 2015; Cropper et al., 2016). In our experiments, for example, CCAP prolonged the protraction phase of the MCN1-gastric mill rhythm while it shortened the retraction phase of the CabPK-gastric mill rhythm. CCAP prolongs protraction during the MCN1-rhythm by activating I_MI_ in the LG neuron which, unlike the parallel MCN1-activated I_MI_ in LG, is not regulated by feedback inhibition (DeLong et al., 2009b).

Despite also sharing with CabPK the ability to activate I_MI_ in LG, CCAP did not alter the LG burst characteristics during the CabPK-gastric mill rhythm but instead decreased its interburst interval (**Fig. 8B**). It was therefore possible that this CCAP effect on the CabPK-rhythm was not due to its action on LG but instead resulted from its ability to increase the pyloric cycle frequency, insofar as the latter action is known to increase the speed of the CabPK-rhythm (Weimann et al., 1997; Saideman et al., 2007a). In our experiments, however, this potential mechanism was unlikely because CCAP did not change the pyloric rhythm speed, presumably because the baseline pyloric cycle frequency was often faster than the limited range in which CCAP increases its speed (≤ 0.5 Hz; Weimann et al., 1997).

Alternatively, the CCAP-activated I_MI_ in LG may influence the CabPK-gastric mill rhythm by summing with the CabPK-activated I_MI_ during the retraction phase. The summed I_MI_ could enable LG to reach burst threshold sooner, possibly by reducing the amount of the second CabPK-activated current (I_Trans-LTS_) that is required to produce the LG burst and/or enabling I_Trans-LTS_ to be activated sooner (Rodriguez et al., 2013). Although the mechanism underlying the CCAP-mediated shortening of CabPK-retraction duration is not established, computational modeling (Rodriguez, 2014) and dynamic clamp experiments (this paper) support the hypothesis that the CCAP influence on the CabPK-gastric mill rhythm does result from CCAP activation of I_MI_ in LG.

In some preparations (13%), CCAP did not alter the CabPK-gastric mill rhythm despite consistently prolonging the MCN1-gastric mill protraction phase. This discrepancy may have resulted from an occlusion effect, insofar as CabPK and CCAP both activate I_MI_ in the LG neuron and their receptors may influence the same pool of I_MI_ channels (Swensen and Marder, 2000; DeLong et al., 2009b; Garcia et al., 2015). If so, then in preparations where the CabPK concentration activated all I_MI_ channels, addition of CCAP would have no effect. Such an occlusion might have occurred in some preparations because the same STG neurons in different individuals express a different maximal conductance for each of its currents (Schulz et al., 2006; Schulz et al., 2007), a feature also recognized in other systems (Luo et al., 1999; Kamme et al., 2003; Cristobal et al., 2005; Swensen and Bean, 2005; Tietjen et al., 2005). This individual variation can result in the same modulatory action having a different impact on circuit output in different preparations (Weimann et al., 1997; Schulz et al., 2006; Spitzer et al., 2008; Gabriel et al., 2009; Grashow et al., 2009; Sakurai and Katz, 2009; Grashow et al., 2010; Marder et al., 2015; Blitz et al., 2019). An occlusion effect would be less likely to occur during the MCN1-rhythm, despite the fact that MCN1-released CabTRP Ia also activates I_MI_ in LG, because MCN1-activated I_MI_ amplitude declines during the protraction phase, whereas CCAP-activated I_MI_ amplitude persists (DeLong et al., 2009b).

The distinct CCAP-mediated changes in phase durations during the CabPK- and MCN1-gastric mill rhythms suggest that there would be different behavioral outcomes for these two rhythms. For example, with respect to the LG neuron, which innervates protractor muscles, selectively altering the LG neuron burst or interburst characteristics would likely alter the duration and amplitude of tension generated in those muscles, thereby changing the chewing pattern (Weimann et al., 1991; Stein et al., 2006; Diehl et al., 2013). Because CabPK is located in a distinct set of projection neurons from MCN1 (Saideman et al., 2007b), it is possible that distinct inputs elicit either CabPK- or MCN1-driven gastric mill rhythms, therefore allowing the same circulating hormone (CCAP) to either reduce retraction duration (CabPK-rhythm) or increase protraction duration (MCN1-rhythm).

### Convergent output can occur through distinct synaptic pathways

GPR stimulation only affects the MCN1 (10-15 Hz)-gastric mill retraction phase (Beenhakker et al., 2005; DeLong et al., 2009a). This action results from the GPR-mediated selective inhibition of MCN1_STG_ neuropeptide release, which prolongs retraction by slowing MCN1 excitation of LG (DeLong et al., 2009a). Stimulating GPR during protraction did not alter either phase, presumably because LG is already inhibiting MCN1_STG_ during protraction (DeLong et al., 2009a,b). Here we showed that GPR stimulation during retraction also prolonged that phase, while stimulating it during protraction reduced protraction duration, during both the MCN1(3-6 Hz)- and CabPK-gastric mill rhythms.

Despite these two gastric mill circuits exhibiting the same response to GPR stimulation, GPR is likely acting via different synapses during the two gastric mill rhythms because MCN1 is either silent or only weakly active during the CabPK-rhythm (Saideman et al., 2007b; this paper). GPR also directly excites Int1 and DG, and directly inhibits LG (Beenhakker et al., 2005; DeLong et al., 2009a) (this paper). While these latter synapses do not mediate the GPR action on the MCN1-rhythm, one or more of them may be responsible for GPR prolonging the CabPK-gastric mill retraction phase. For example, GPR excitation of Int1 and/or DG activity could strengthen their inhibition of LG during CabPK-retraction, potentially slowing the ability of LG to generate its next burst, as would direct GPR inhibition of LG. We eliminated DG as being necessary for this action, but the relative influence of the GPR actions on Int1 and LG remains to be determined. This convergent action of GPR on the distinct MCN1- and CabPK-gastric mill circuit states, albeit via different GPR synapses, also suggests that this feedback pathway plays a pivotal role in producing a particular chewing behavior *in vivo*, regardless of the underlying circuit state. Such a convergence is not the only possible outcome, as different circuit states can also respond differently to consistent sensory activation (Gordus et al., 2015).

Circuit flexibility is not unique to the STNS. For example, the same intersegmental circuit generating the leech heartbeat rhythm regularly switches to produce synchronous and metachronal patterns (Calabrese et al., 2016). Additionally, the swimming circuit in two nudibranch mollusk species is composed of the same set of neurons using different circuit configurations (Sakurai and Katz, 2017), and different circuit configurations distinguish the gastric mill circuit in *C. borealis* and the lobster *Panulirus interruptus* (Elson and Selverston, 1992; Nusbaum et al., 2017). Activating different inputs can also configure different phase relationships among feeding circuit neurons in the molluscs *Aplysia californica* and *Lymnaea stagnalis*, producing ingestion or egestion (Cropper et al., 2016; Crossley et al., 2018), and in *Aplysia*, egestion can be generated by two degenerate circuit states (Wang et al., 2019). In the latter case, the functional consequence is a substantial difference in the latency for switching from generating the egestion motor pattern to that for ingestion in response to an ingestion-activating signal.

In summary, our data extend the known functional consequences of degenerate circuit states by showing that, while in some cases such circuit states do respond differently to the same external influence, they can also respond similarly to the same extrinsic input, albeit not necessarily via the same synaptic pathway. This convergent response to the same input increases the challenges associated with interpreting comparable results in less well-defined systems where a consistent behavioral or network response to a perturbation occurs across individuals or at different times in the same individual.

## Acknowledgments

Dr. Jason C. Rodriguez performed preliminary experiments that led to several of the hypotheses addressed here. Dr. Ekaterina Morozova provided essential guidance pertaining to dynamic clamp experiments.

